# An ethogram method for the analysis of human distress in the aftermath of public conflicts

**DOI:** 10.1101/2023.05.30.542823

**Authors:** Virginia Pallante, Peter Ejbye-Ernst, Marie Rosenkrantz Lindegaard

**Affiliations:** Netherlands Institute for the Study of Crime and Law Enforcement (NSCR), Amsterdam, The Netherlands; Department of Sociology, University of Amsterdam, Amsterdam, The Netherlands; Department of Sociology, University of Copenhagen, Copenhagen, Denmark

**Keywords:** distress, post-conflict, human behaviour, real-life observation, interdisciplinary approach

## Abstract

Research on nonhuman animals has widely documented the behavioural expression of distress in a conflict context. In humans, however, this remains largely unknown due to the lack of direct access to real-life conflict events. Here, we took the aftermath of 129 video recorded street fights and applied the ethological method to explore the behavioural cues of people previously involved in a fight. Drawing on observations on nonhuman behaviour and inductively identified behaviours, we developed and inter-coder reliability tested an ethogram for the behavioural repertoire of distress. We further quantitively analysed the behaviours with a correlation matrix and PCA, that revealed that the behaviours we observed were not displayed in combination with each other, showing a variability in how people express distress. Since both human and nonhuman primates react to conflict situations with similar expressions of distress, we suggest a comparative approach to understand the evolutionary roots of human behaviour.

## 1. INTRODUCTION

Decades of ethological research on nonhuman animals have explored the social dynamics occurring in conflict events, investigating the role of emotions in affecting a wide range of resolution and management strategies aimed at maintaining social cohesion (Aureli & de Waal, 2000). The occurrences of these reparative mechanisms are largely affected by the emotional state of the previous antagonists, which regulate their subsequent interactions and bystanders’ propensity to intervene (Clay & de Waal, 2013; Heesen et al., 2022). Although conflict reparative strategies have been documented recently in humans (Lindegaard et al., 2017; Philpot et al., 2022), how emotions are expressed by the people who took part in the conflict affect conflicts’ dynamics remains unexplored. As for nonhuman primates, it is expected that in a conflict context humans experience a state of distress (Fujisawa et al, 2005). However, because studying distress in real life is challenging, its research has typically been confined to controlled artificial settings, mostly relying on physiological reactions and psychological measures, the latter being frequently questioned due to the cognitive biases or inaccurate self-evaluation of participants (Troisi, 2002). We might ask, though, whether the behavioural cues that laboratory studies have documented as the expressions of emotions are actually generalizable to real-life contexts.

With the aim to advance the understanding of the role of emotions in shaping conflict dynamics in humans, we explored the behavioural repertoire for the expression of emotions displayed by people who have just taken part in street fights in public spaces. Our purpose with this analysis was to provide a tool to detect and classify potential behavioural indicators of distress in the tense context that follows a conflict event, and to systematically study this behavioural repertoire in naturally occurring situations.

Although both in human and nonhuman animals a window on the emotional state is provided by the spontaneous behaviours that an individual displays (Ekman 1965, 1998; Argyle, 2013; Darwin, 2009[1872]) only a few studies have focused on the behavioural repertoire of people as a response modality to distress (Troisi, 1999, 2002; Sgoifo et al., 2003; Bardi et al. 2011; Whitehouse et al., 2022). Drawing from observations on nonhuman primates, this body of research has, for example, documented displacement activities as effective behavioural indicators of distress in humans, supporting the validity of behavioural observations as a complement to physiological and psychological measurements as a way to measure a state of distress and suggesting a common root with other nonhuman primate species in the evolution of the behavioural response to distress (Maestripieri, 1992; Troisi, 1999, 2002, Whitehouse et al., 2022). In particular, displacement activities have been identified as indicators of anxiety, an affective state characterized by tension and/or agitation. In humans, they include self-directed actions, such as touching the face or passing the fingers through the hair in a combing movement, and aimless manipulation of objects, for example twisting and fiddling finger movements with small items (Troisi, 1999, 2002). Repetitive actions in fixed forms with no obvious purpose are also displayed to discharge tension and anxiety (Troisi, 2002). Moreover, it has been documented that social stress elicits subsequent aggressive interactions, known as displacement aggression, which have been interpreted as reactions to dissipate anger (Fitz, 1976; Marcus-Newhall et al., 2000). The first systematic behavioural observations have been conducted on patients suffering of chronic distress due to mental disorders (Troisi 1996, 1999). The subsequent studies on non-clinical subjects have relied on the findings of this research rather than being explorative (Sgoifo et al., 2003; Pico-Alfonso et al., 2007; Mohiyeddini & Semple, 2013; Zandara et al., 2018). As a result, we do not know whether the behavioural repertoire elicited via standardized protocols inducing mild distress in the participants in such artificial settings and for individuals who do not suffer from chronic distress due to mental disorders presents a wider variability, nor whether it is ecologically valid in situations encountered in real-life interactions.

To answer these questions, we need to investigate the expression of distress in a real-life context. Naturalistic observations of spontaneous behaviours displayed in tense situations are limited to few examples that include waiting rooms of dental offices (Barash, 1974) or medical practices (Shreve et al., 1988). In these cases, the relation between the behaviours observed and the feeling of distress underpinning them is merely assumed by the context where they occur, that in the eyes of the researcher is evaluated as stressful.

A less explored context characterized by tension is the period that follows an interpersonal conflict. Although behavioural studies of emotions in humans in such situation are virtually absent, research on nonhuman animals highlight that the post-conflict context is characterized by high levels of anxiety (Aureli & van Schaik, 1991; Castles & Whiten, 1998; Das et al., 1998; Kutsukake & Castles, 2001; Palagi et al., 2014). Observations on nonhuman primates documented that the subjects who were involved in a previous fight increase their expression of displacement activities such as scratching, self-grooming, and body shaking (Maestripieri, 1992; Schino et al. 1996; Bardi et al. 2003, 2004; Leavens et al. 2004; Kalueff et al. 2010). Furthermore, subjects in the aftermath of a conflict might reiterate their aggressive motivation against other group members (Romero et al., 2011; Pallante et al, 2018). Similar findings have been obtained in field studies on children. For instance, after being involved in an aggressive interaction with peers, children show displacement activities, anger and sadness as expression of distress (Arsenio & Killen, 1996; Butovskaya & Kozintsev, 1999; Ljungberg et al., 1999; Fujisawa et al., 2005; Westlund et al., 2008). Understanding the behavioural cues of distress in human adults in the aftermath of an interpersonal conflict would therefore contribute to characterize the post-conflict as a context of tension and add the expression of distress as one of the potential factors affecting the subsequent social dynamics, including for example bystanders’ decision to intervene or provide consolation (Lindegaard et al., 2017). It would further contribute to highlight the phylogenetic proximity of humans with other nonhuman primate species in the expression of distress.

In order to explore the behavioural repertoire of people as a response modality to distress in real-life tense events and given the importance of understanding the role of distress in naturally occurring aggressive interactions, we take the post-conflict context as our object of investigation assuming it is a context of tension. By conducting a systematic video observation of people who were involved in a street fight, the aim of this study was to establish how people express behavioural cues of distress. We also aimed to test the reliability of this classification tool for the systematic study of distress. For this purpose, we adopted the ethological method that researchers developed for the study of animal behaviour (Altman 1974; de Waal & Yoshihara, 1983; Lehner 1998).

A valid tool recently adopted in criminology to access real-life contexts consists of the footage recorded by Close Circuit Television (CCTV), which offers direct observation to crimes occurring in public spaces (Lindegaard & Bernasco, 2018; Philpot et al., 2019). CCTV footage allow researchers to explore the behavioural repertoire displayed during events such as street fights (Levine et al., 2011; Ejbye-Ernst et al., 2020). Scholars have used CCTV footage to evaluate the role of emotions on the outcomes of robberies by inferring the emotional state of victims and robbers from the behaviours they display (Mosselmann et al., 2018; Nassauer, 2018). In documenting these behaviours, the authors associated the expression of emotions to a specific body language, including facial expressions, body movements, and postures. A similar approach has been used to describe the behaviours leaking information about emotions of people involved in violent events and their effect on the escalation of violence (Klusemann, 2009; Nassauer, 2016). These studies qualitatively approached the behavioural expression of emotions in naturally occurring aggressive interactions, highlighting their communicative value in the social dynamics of the events. Although the ethological observational method inspired the systematic quantitative approach in the study of CCTV material (Lindegaard & Bernasco, 2018), no attempt to apply it for the detection of the expression of the distress has been made yet.

In this paper, first, we developed an ethogram through inductive observation of real-life conflicts where we detailed a qualitative description of the behaviours that potentially indicate a distressed condition in the post-conflict context. Before the possibility to video observe people in real-life contexts, ethograms were rarely developed for the study of human behaviour (Eibl-Eibesfeldt, 2007; Jones et al., 2016), although they have proved to be a valid tool to detect the behavioural indicators of distress in clinical research (Troisi, 1999, 2002). Second, we tested the reliability of the ethogram with two independent observers. Third, we showed how to use the ethogram by systematically recording and quantifying the behaviours observed, providing an overview of how these behaviours are expressed in the post-conflict context. Our work aimed to offer other researchers a toolkit to assess the behavioural expression of distress in conflict dynamics and its potential influence on the consequences of violence. By conducting a behavioural analysis, it will further allow for a comparative approach in the study of distress that has the potential to highlight a phylogenetic continuity of humans with other nonhuman primate species.

## 2. MATERIALS AND METHODS

### 2.1. Participants

We analysed data from a sample of CCTV footage recorded by the police of Amsterdam. We gained permission to use the footage for scientific purposes from the Dutch Prosecution Services. The footage came from cameras located in the city of Amsterdam (The Netherlands) and were recorded by municipal camera operators from March to June 2020. Footage was recorded during the COVID-19 pandemic and involved incidences observed by operators as part of their daily routines. The conflicts sampled included varying levels of intensity, and for each conflict 30 min before and after the event were recorded. Therefore, each clip had a duration of approximately one hour. The footage did not have sound.

Out of a total sample of 1112 clips, 65 involved clear conflicts between citizens. The conflict parties were all adults, except for 12 episodes of bullying that involved adolescents. Additionally, we included 11 videos in our sample where no conflicts were visible, but the operators started the recording as soon as the conflict had ended (within 10 min). In these cases, we had information about the content and the characteristics of the preceding conflict because the operators write a brief description of these missed episodes. The final sample consisted of 76 clips that met the following criteria:

1. The quality of the video (resolution and brightness) was high enough to allow for coding.
2. At least one antagonist was present in the post-conflict period.
3. The antagonist was clearly visible for at least 1 min after the end of the conflict.
4. In our total sample, we conducted observations on 126 antagonists, of which 11 were women and 115 were men.

### 2.2. Observation and coding procedures

Post-conflict observations were conducted with the intent to (1) develop the ethogram—in a first observation phase, and (2) code and quantify the behaviours displayed—in a subsequent coding phase.

#### 2.2.1. Observation phase

We focused our observations on the antagonists of the conflict, i.e., the persons who were directly involved in the immediately preceding fight. The conflict parties were labelled with the neutral term of antagonists or opponents instead of actor and receiver of violence, since within a single conflict they might equally display and receive violence. Bystanders who intervened in the conflict were excluded from the study. Observations were conducted in the post-conflict period, i.e., as soon as the conflict was over. We defined the end of a conflict as when the antagonists have stopped to aggressively interact with each other, i.e., they do not show any threatening or aggressive behaviours, and have moved their attention away from their previous opponent. In the 11 cases where the conflict was not recorded, observations started as soon as the antagonists were visible in the frame. During the post-conflict phase, the antagonists occasionally remained in the same location and engaged in subsequent shorter, less intense, aggressive interactions (i.e., characterized by the absence of physical contact and shorter compared to the previous aggression). In these cases, we considered the most intense aggression as the main conflict and coded the subsequent less intense episodes as redirected aggression expressing distress.

In order to develop the ethogram, we followed two different procedures. First, a deductive phase consisted in identifying the presence of the behaviours related to distress that were already documented in the existing literature (Troisi, 1999, 2002). However, since the relevant literature focused on distress in contexts different than the post-conflict one, we adopted a flexible approach for the identification of the behavioural patterns, given that they could slightly change in their expression according to the situation. Therefore, in order to encompass this behavioural variability and to identify additional behaviours displayed by the antagonists, we combined the deductive phase with an inductive approach by conducting *ad libitum* observations (Altmann, 1974). During this second phase, we recorded all the most frequently observed behaviours in order to understand whether their occurrence might characterize the post-conflict context. These behaviours consisted of behavioural patterns that were clearly discernible, limited, and repeated over time, including both individual activities and interactions between antagonists and third parties.

All the behaviours recorded in the inductive and deductive phases were labelled as states or events. The previous literature defined a state as any ongoing behaviour characterized by a relatively long-lasting duration, for example a prolonged activity or a posture. Conversely, an event has been described as an instantaneous occurrence or momentary behaviour, such as a precise body movement (Altman 1974; de Waal & Yoshihara, 1983; Lehner 1998). Following this distinction, we categorized as states prolonged social interactions, individual activities and body postures, and as events instantaneous movements and gestures of a negligible duration (e.g., a single self-touching movement). We provided the distinction between states and events both to define the characteristics of the behaviours observed in terms of duration and to propose an operative method for their coding (see the section Coding phase). The observation phase ended as soon as both antagonists were out of frame. A total of 1305 min was observed, with a mean duration per person of 10.61 min, a minimum of 1 min, and a maximum of 38 min per person.

#### 2.2.2. Coding phase

We conducted a systematic encoding of the CCTV material to quantify the behaviours observed in the first phase by applying the focal sampling technique (Altmann, 1974). Using the ethogram developed in the observation phase, we coded the behaviours of one antagonist at a time for at least one minute and up to (and no more than) 10 min after the end of the conflict. We stopped the coding if police officers arrived and handcuffed or arrested the antagonist, in order to limit his/her actions. Similarly, observations ended when the personnel of an ambulance provided help to an injured antagonist constraining his/her possibility to move. A total of 872 min was coded, with a mean duration per person of 7.09 min.

During the coding phase, the behaviours were coded per minute. For every minute of observation, the coder recorded all the behaviours an antagonist displayed, the time of their occurrence, and the receiver of the behaviour in cases of interactions.

Behaviours were coded according to their definition as state or event. An event was coded every time it was observed, irrespectively from its duration, while a state was coded only when it lasted at least 2 s without any interruption. Two events of the same type (e.g., self-touching) were coded twice when they represented different forms of the same behaviour (e.g., touching the face and touching the hair, included into the same category in the ethogram), or they occurred twice but they were separated by at least one second (e.g., touching the face and after one second touching again the face). Two states of the same typology were recoded twice if they were separated by at least 2 s, otherwise they were counted as a single occurrence. In the case of interactions directed at someone (e.g., talking with gestures and talking with aggressive gestures), the same behaviour was coded twice if the receiver changed during the interaction. Although this does not pose any problem when coding events, in order to capture the occurrence of a state that started in a given minute and continues in the following minutes, we reported the coding of the same state again at the beginning of each minute. When the focal person moved out of the frame, we stopped coding only if he/she was not visible for at least 10 s continuously, and resumed the coding when the person was back in the frame. Shorter out of sight behaviours were not considered. All the behaviours were manually coded in an excel sheet.

### 2.3. Statistical analysis

In order to explore the variability in the expression of distress and to understand whether people express specific behaviours in combination with each other, we ran a correlation matrix with the frequencies of the behaviours. In addition, we ran a Principal Component Analysis (PCA, no rotation) on the matrix of the behavioural frequencies to evaluate whether the behaviours observed could be empirically aggregated in categories to explain their variance in the post-conflict context. With the PCA, we aimed at understanding if behavioural patterns exist in the way people express a state of distress. We opted for an empirical analysis to reduce the complexity in the expression of the behaviours observed and since no previous theoretical background exists assuming any a priori potential aggregation of the behaviours indicating distress. All the statistical analysis were run in R.

### 2.3. Ethical statement

The study was conducted in accordance with the Declaration of Helsinki. The project has been approved by the Netherlands Public Prosecution Service (PaG/BJZ/49986) and assessed by the Ethics Review Board of the Faculty of Social and Behavioral Science at the University of Am-sterdam (2022-AISSR-15534).

## 3. RESULTS

### 3.1. The ethogram

The ethogram consists of 22 different behaviours, including 9 events and 13 states (Table 1). Since the co-presence of states and events might result in a heterogenous ethogram, we grouped all the behaviours according to their shared characteristics, which further emphasize their dynamicity: Self-directed behaviours, Body postures, Movements, Interactions with objects, and Interactions with people. In the ethogram, each behaviour is followed by a short code to facilitate its subsequent recording, and by a qualitative description. The behaviours recorded consisted of gestures, postures, interactions, and movements. Since the quality of our material did not always allow to clearly identify facial expressions, we included their occurrence in combination with some behaviours, but we avoided their recording as self-standing patterns and consequently their coding. We attributed the same behavioural code to subtle different forms of the same behaviour, in order to preserve the variability observed by considering all the patterns expressed, but at the same time to avoid a too detailed coding of similar behavioural patterns. Self-touching, for example, could be expressed by touching different parts of the body (hair, face, arms, etc.). Although we reported these different forms in the description of self-touching in the observation phase, in the coding phase we coded each variation in this behaviour simply as Self-touching.

**Table 1.**
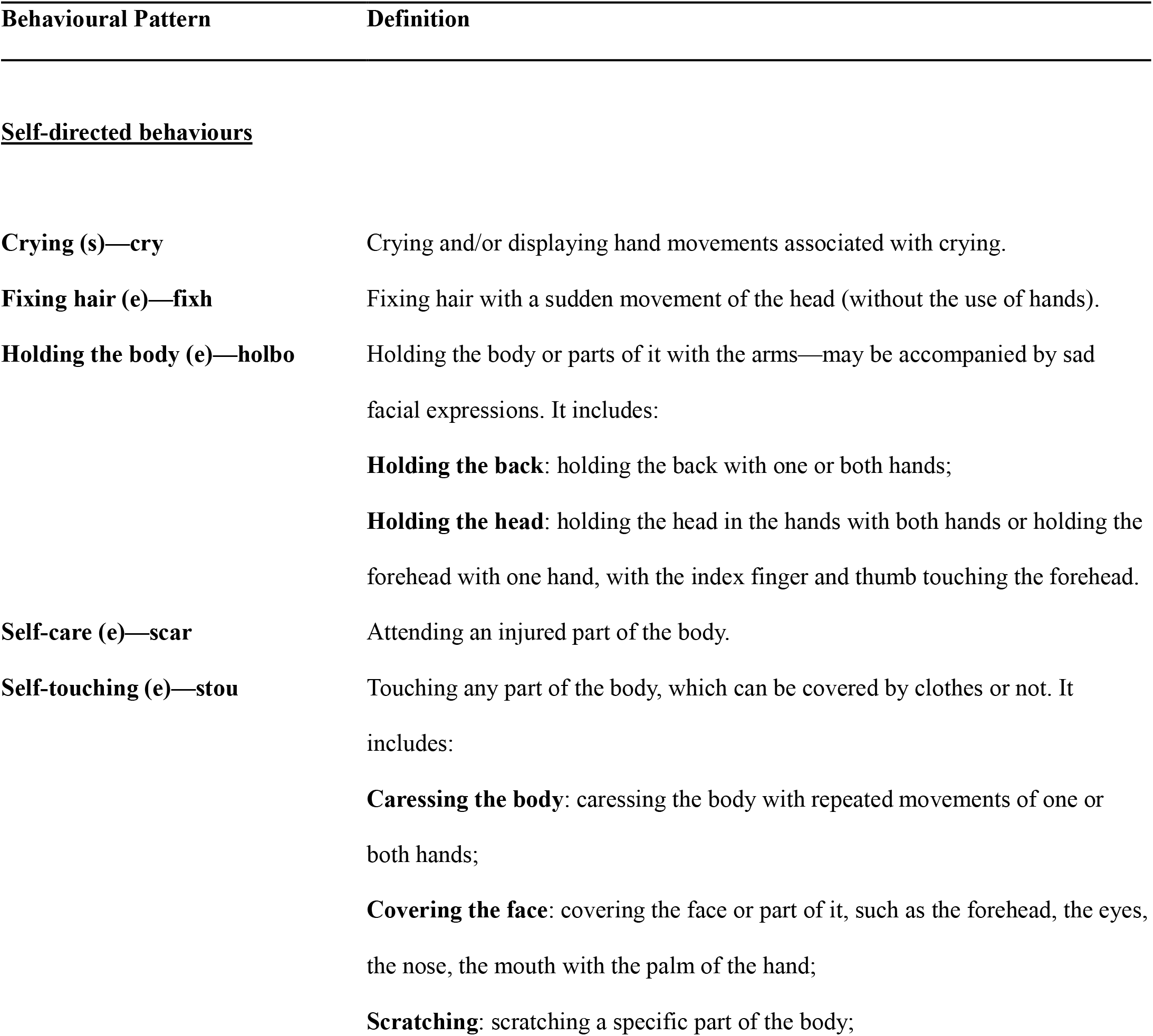
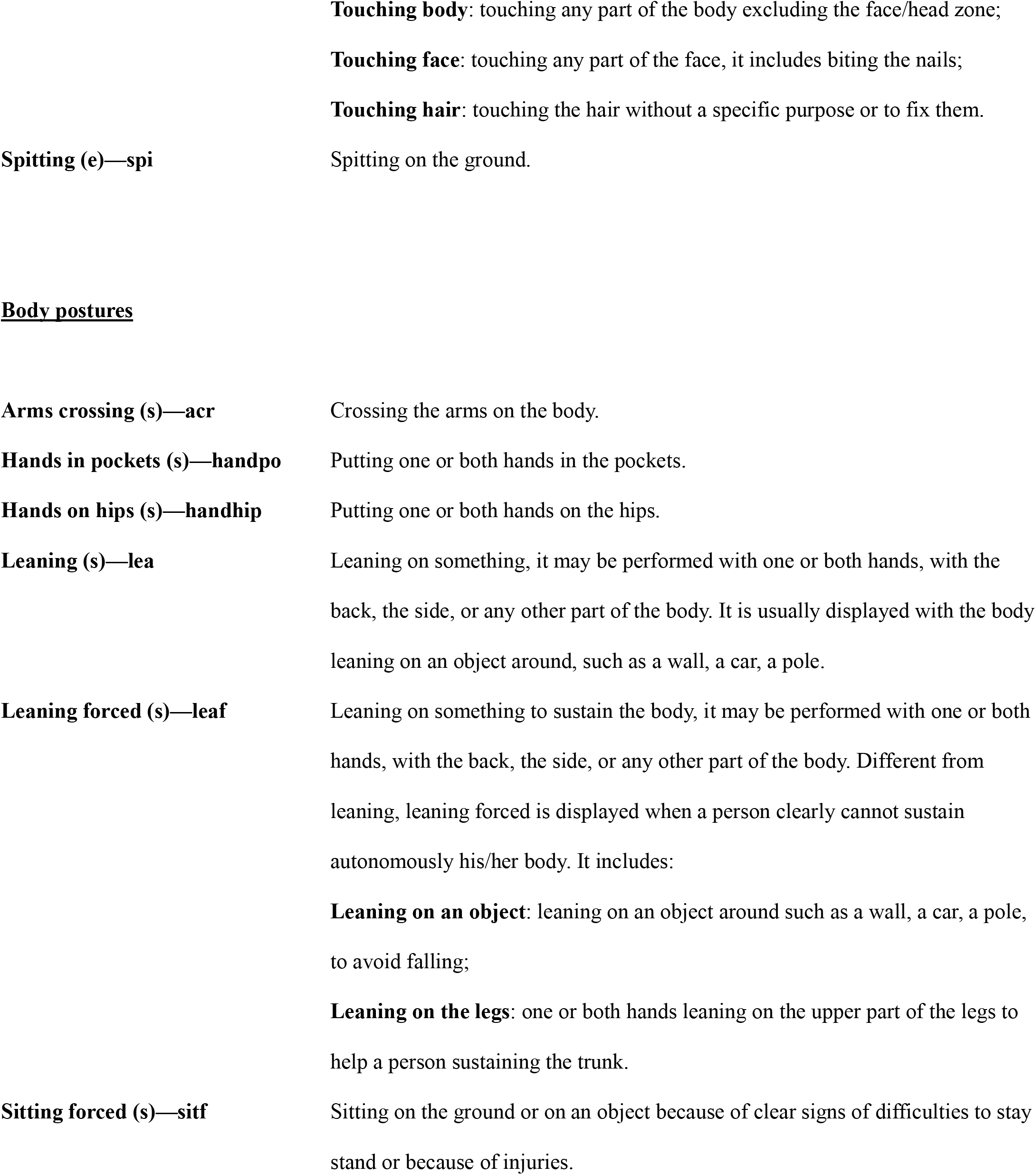
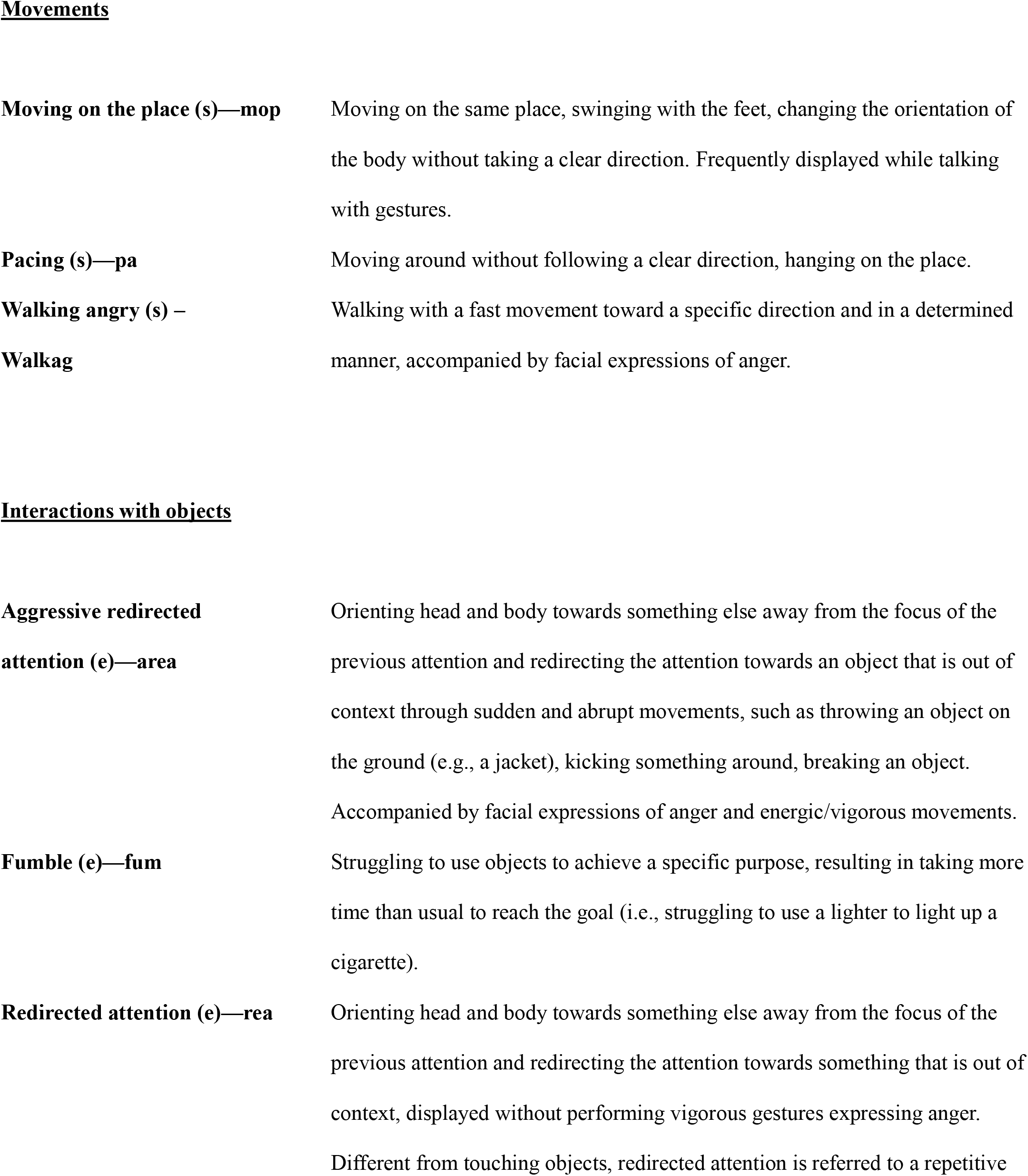
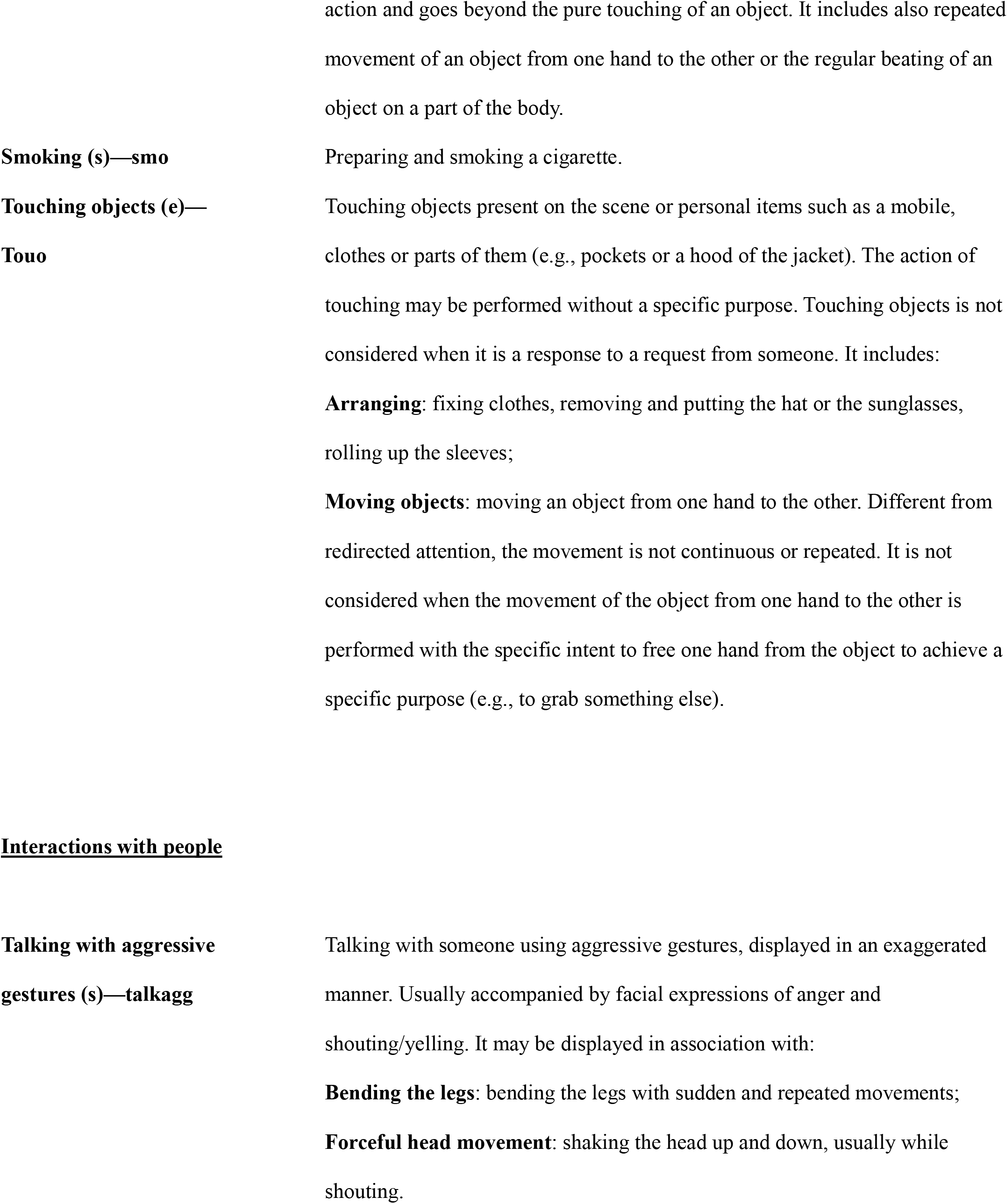
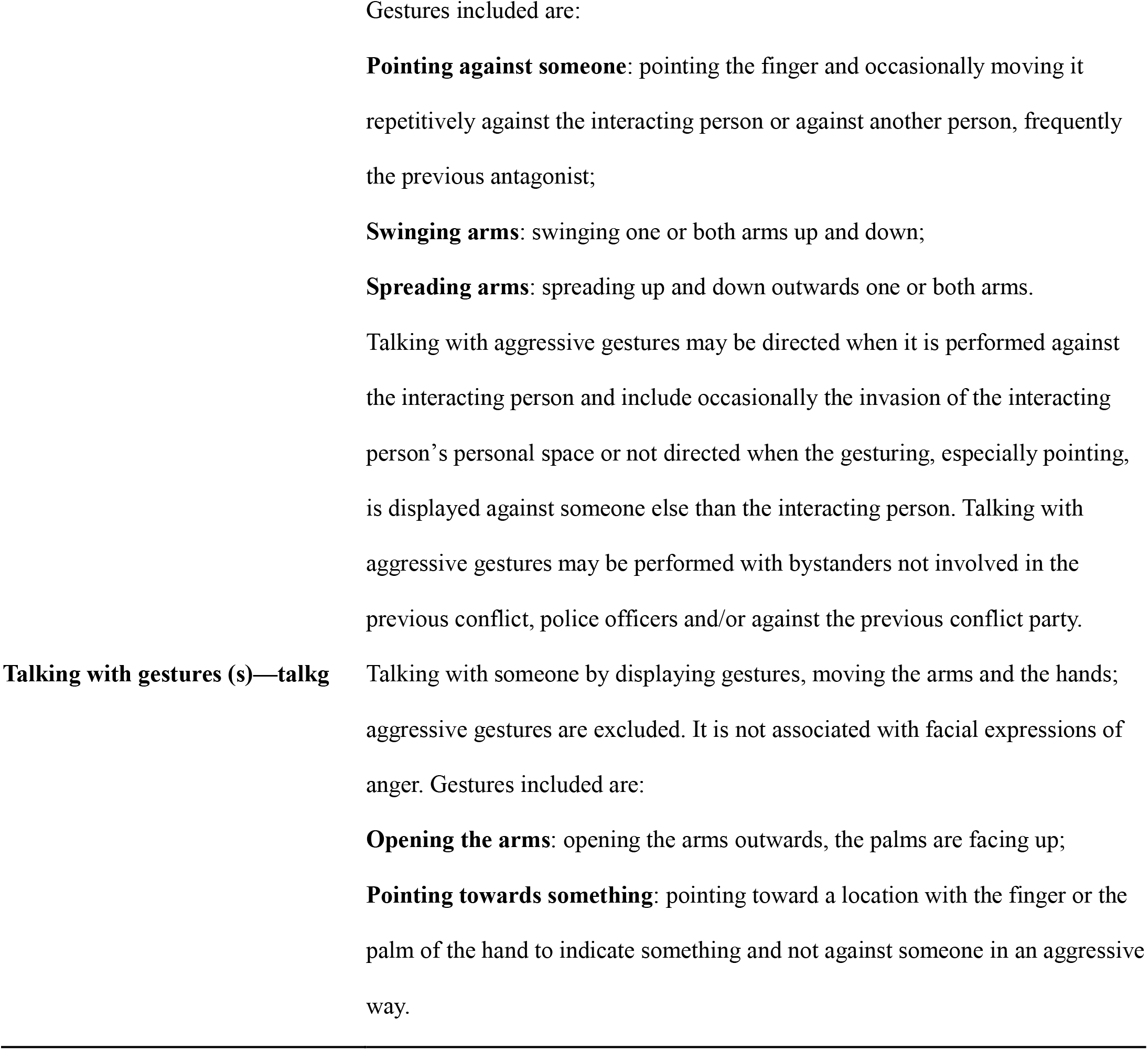
Ethogram and definitions of the behaviours observed. The behaviours are identified as states (s) or events (e).

We further combined the observed behaviours into three categories according to their characteristics and their possible relation with the underpinning emotional state of distress (Table 2).

**Table 2.** Behavioural patterns grouped per category.

| CATEGORY 1 | CATEGORY 2 | CATEGORY 3 | CATEGORY 4 |
| --- | --- | --- | --- |
| Anxiety | Anger | Physical Sufferance | Unclassified Behaviors |
| Fixing hairs | Aggressive<br>redirecting attention | Crying | Arms crossed |
| Fumble | Talking with<br>aggressive gestures | Leaning forced | Hands in pockets |
| Holding the body | Walking angry | Self-care | Hands on hips |
| Moving on the place |  | Sitting forced | Leaning |
| Pacing |  |  | Spitting |
| Redirecting attention |  |  | Talking with gestures |
| Self-touching |  |  |  |
| Smoking |  |  |  |
| Touching objects |  |  |  |

Category 1 included self-directed behaviours such as “Self-touching” (Figure 1), “Holding the body“ (Figure 2), “Fumble” (Figure 3), and “Touching objects”.

We included in Category 2 aggressive interactions that the antagonists engaged in with other people and that were characterized by gestures expressing anger displayed during a verbal interaction (“Talking with aggressive gestures”; Figure 4).

Category 3 refers to the behaviours that indicate physical sufferance and the self-directed behaviours aimed at attending injures due to the previous aggression (Figure 1). We further included body postures that help a person to sustain the body. Differently from the previous categories, Category 4 includes the behaviours that do not refer to any emotional state but that nonetheless we frequently observed occurring in the post-conflict context. They include body postures and gestural talking (Figure 3).

**Figure 1.**
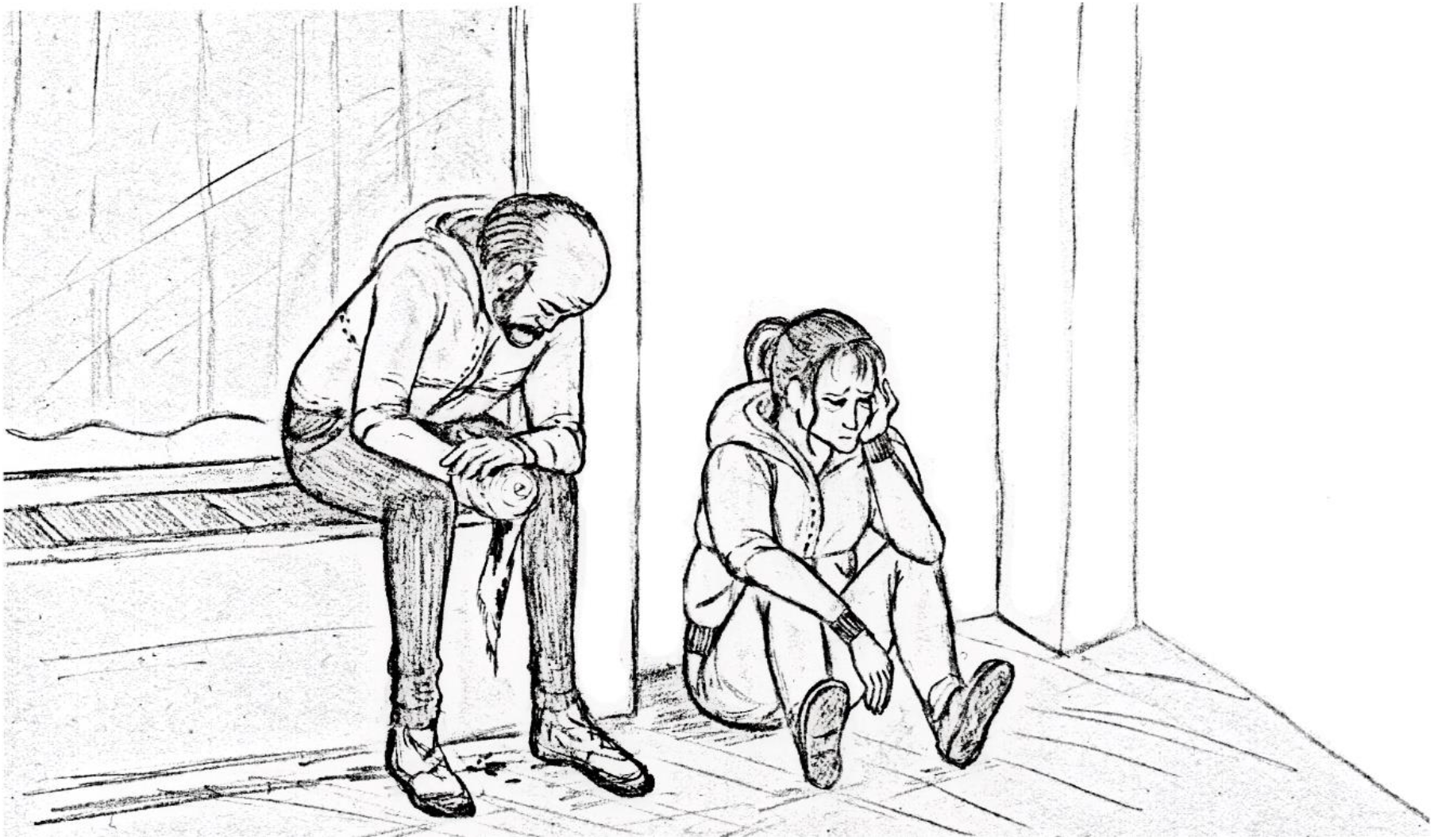
On the right, a girl is touching her face (Self-touching, Category 1) and she sits on the ground (Sitting forced, Category 3) after showing difficulties in standing. On the left and close to her, a man attends his injured hand (Self-care, Category 3), while he sits on a windowsill (Sitting forced, Category 3) leaning his arms on the legs (Leaning forced, Category 3).

**Figure 2.**
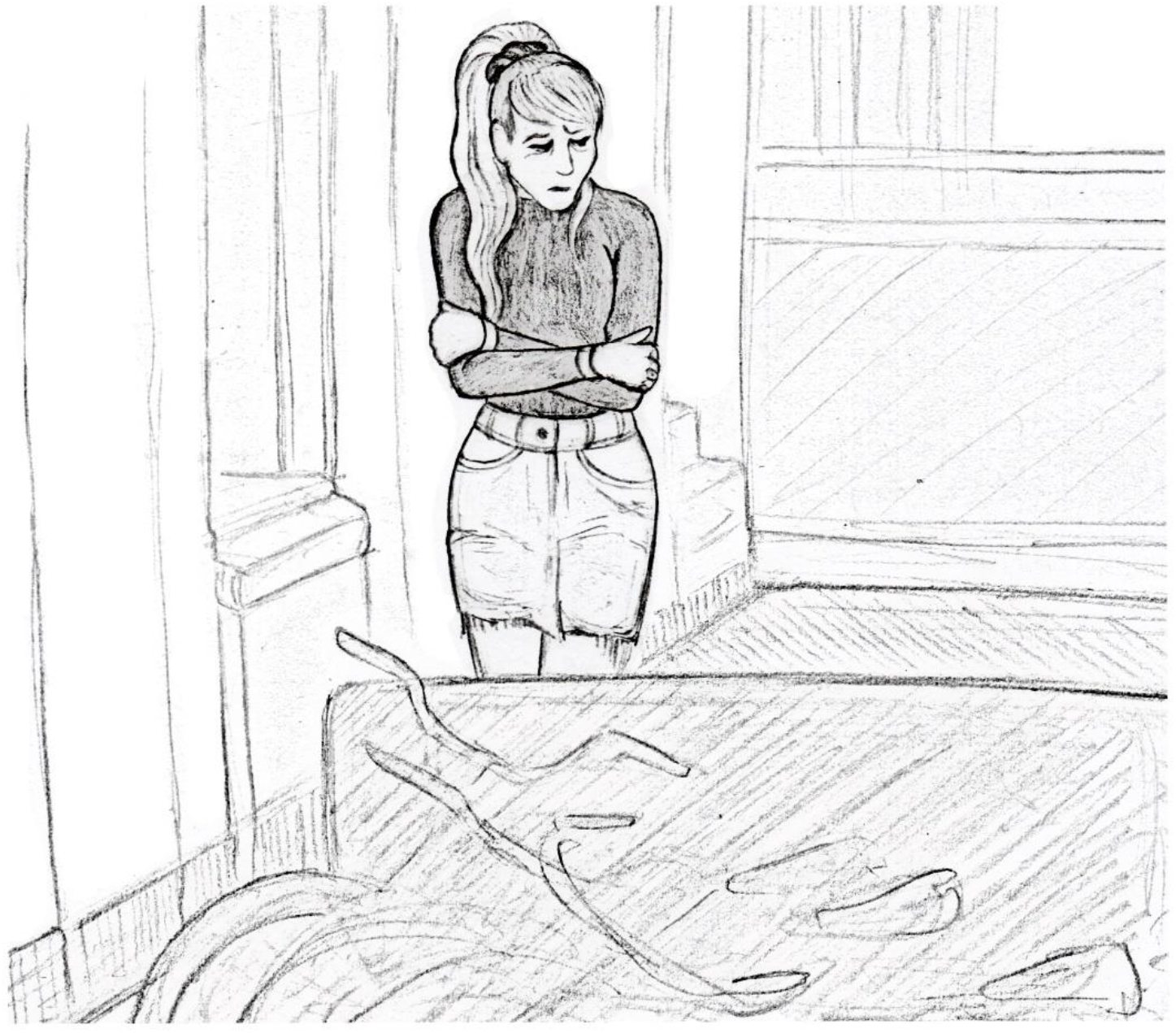
A girl holds her body (Holding the body, Category 1).

**Figure 3.**
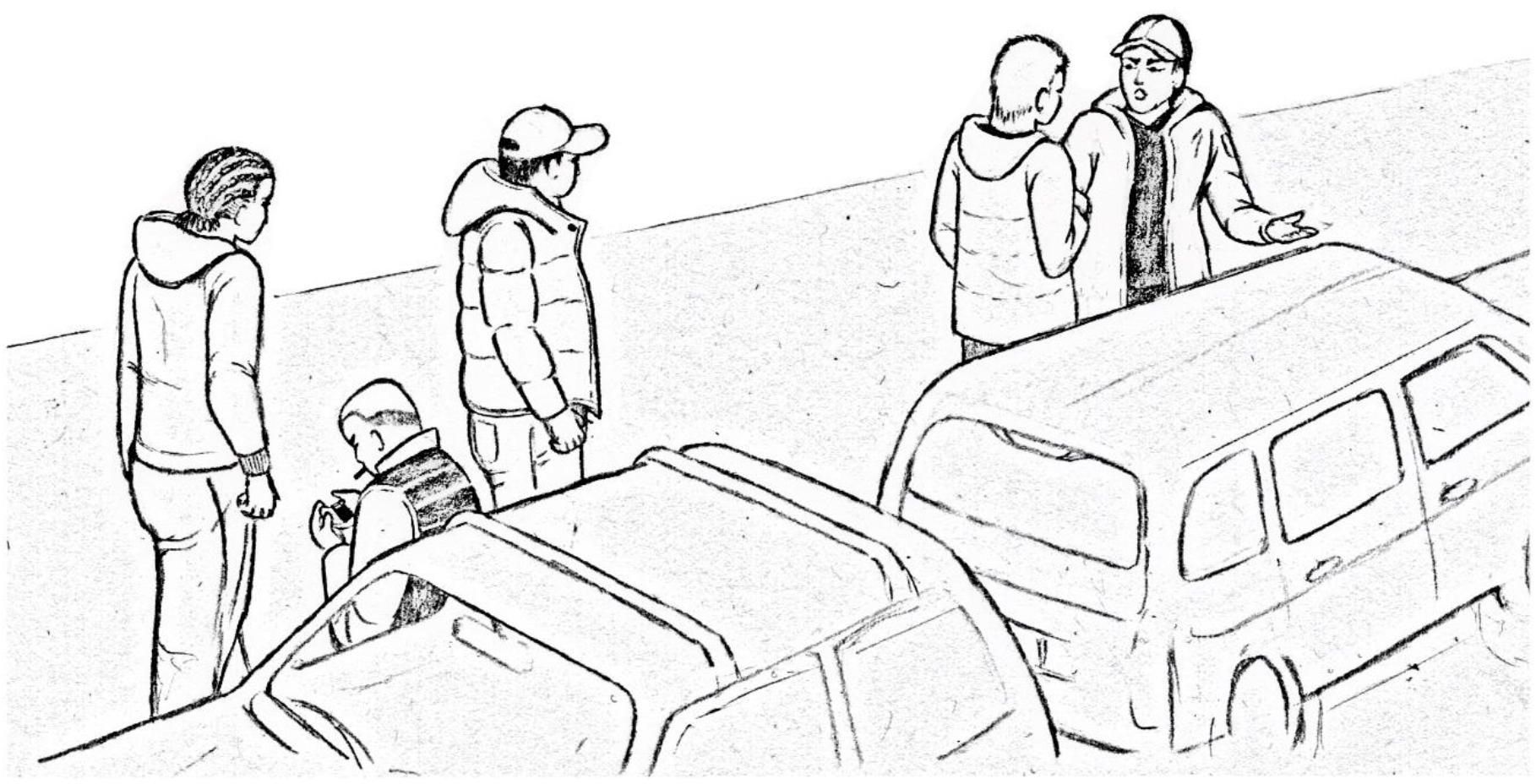
A boy on the left side of the picture is fumbling with a lighter (Fumble, Category 1) while leaning on a car (Leaning, Category 4). He engaged in a physical fight with the boy on the right side, who is talking with gestures with his fellow friends (Talking with gestures, Category 4).

**Figure 4.**
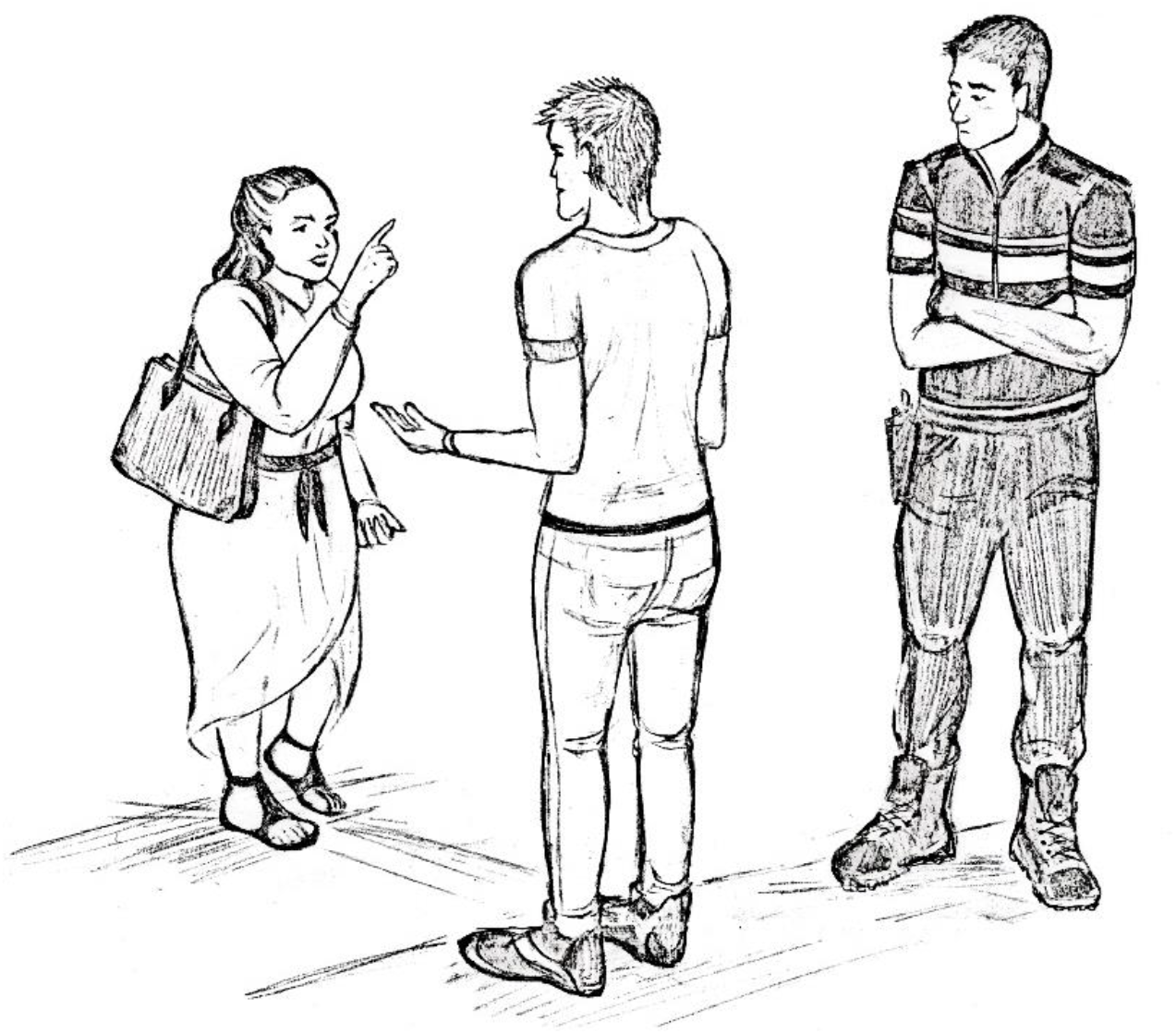
A woman is talking using aggressive gestures (pointing) against a man who was not involved in the previous conflict event (Talking with aggressive gestures, Category 2).

### 3.2. Ethogram validation: assessing the intercoder reliability

After the development of the ethogram and prior to the coding of all the video material, a second independent observer was trained to test the reliability of the ethogram. The agreement was estimated by double-coding and comparing approximately 10% of the entire material, randomly selecting 23 antagonists from the sample. In total, we recorded and compared 387 behaviours.

During the coding phase for testing the intercoder reliability, each coder received a list of codes and recorded in an excel sheet the presence/absence of the behaviour within each minute. The intercoder reliability was estimated through the Cohen’s Kappa (κ). The results are reported in Table S1 and show substantial agreement for all the behaviours (Landis & Koch, 1977).

### 3.3. Quantitative results

Across the entire sample, we coded a total of 3719 behaviours. Table 3 reports the total amount and the frequencies of each behaviour observed. In general, the most frequent behaviour is “Talking with gestures”, which is part of Category 4. It is followed by “Touching objects” and “Self-touching”, both in Category 1. The most frequent behaviour of Category 2 is “Talking with aggressive gestures”, while for Category 3 is “Sitting forced”. “Aggressive redirected attention” was the less frequent behaviour, with only five cases observed in the entire sample.

**Table 3.** Count and frequency of the total behaviours observed. The frequency is calculated dividing the total occurrences of the behaviours on the total minutes observed (872).

| Behaviour | Total Occurrences | Frequency |
| --- | --- | --- |
| Arms crossing | 78 | 0.089 |
| Aggressive redirected attention | 5 | 0.006 |
| Crying | 11 | 0.013 |
| Fixing hair | 12 | 0.014 |
| Fumble | 9 | 0.010 |
| Hands in pockets | 319 | 0.366 |
| Hands on hips | 66 | 0.076 |
| Holding the body | 8 | 0.009 |
| Leaning | 81 | 0.093 |
| Leaning forced | 35 | 0.040 |
| Moving on the place | 295 | 0.338 |
| Pacing | 93 | 0.107 |
| Redirected attention | 54 | 0.062 |
| Self-care | 43 | 0.049 |
| Sitting forced | 50 | 0.057 |
| Smoking | 52 | 0.059 |
| Spitting | 22 | 0.025 |
| Self-touching | 513 | 0.588 |
| Talking with gestures | 795 | 0.912 |
| Touching objects | 727 | 0.834 |
| Talking with aggressive gestures | 329 | 0.377 |
| Walking angry | 35 | 0.040 |

The results of the correlation matrix revealed no systematic co-occurrence between the variables except for “Sitting forced” and “Leaning forced” (Figure S1 and Table S2).

The PCA did not reveal any meaningful co-occurrence of behaviours. For this reason and in order to give an overview of the results, we report here only the first five dimensions extracted, which explained 39.2% of the initial total variance (Figure S2). The percentage of the next dimensions is reported in Figure S2, where the flat curve we obtained suggests that we need several dimensions to explain the whole variance. Correlation values are reported in Table S3 (see also Figure S3) and the eigenvalues in Table S4. The first component, Figure S4 (PC1), explaining 9% of the variance, was mainly defined by the behavioural categories “Armed crossed” and “Pacing” (both positive correlation). The second component (PC2), explaining 8.5% of the variance, was defined by “Leaning forced” and “Sitting forced” (both negative correlation). The third component (PC3) explained 8.2% of the variance and was defined by “Crying”, “Fixing hair”, and “Self-touching” (positive correlation). The fourth (PC4) and fifth components (PC5) explained, respectively, 7.3% and 6.2% of the variance and were defined by “Moving on the place” and “Talking with gestures” for the first case and “Fumble” for the second (all positive correlation).

## 4. DISCUSSION

In this study we have explored human antagonists’ behavioural cues as potential indicators of distress in real-life conflict events. For this purpose, our first aim was to develop an ethogram of the behaviours displayed by the antagonists who took part to inter-personal conflicts and to test its reliability with a second coder. The agreement reached revealed that the observations matched between the two coders, showing that our main findings are based on reliable observations.

The behaviours we observed included body postures, interactions with people or with objects, movements characterizing different ways of walking, and self-directed behaviours. We grouped all the behaviours into four different qualitative categories in relation to the specific emotional state they might indicate (Table 2). The first category (Category 1) was related to the emotional state of anxiety and included displacement activities, such as self-touching events. These behaviours have been previously identified as expression of a state of anxiety both in human and nonhuman primates, thus supporting our interpretation (Maestripieri et al., 1992; Troisi 1999, 2002). Similarly, self-directed behaviours such as gestures aimed at arranging clothes (“Arrange”) and fixing hair (“Fixing”) also indicate anxiety, as they may convey the same meaning of self-grooming patterns (see Troisi 1999). In addition, repetitive locomotory movements (“Moving on the place”, “Pacing”) and actions that the antagonists performed to redirect their attention towards something out of context (“Redirected attention”) can be interpreted as displacement activities (Maestripieri et al., 1992; Troisi 2002). Finally, we included in Category 1 smoking, as field and laboratory studies reported its occurrence as a response to anxiety (Niaura et al., 2002). The behaviours we included in Category 2 share the common characteristic of being an expression of anger, that we identified also through facial expressions and sudden movements which are displayed faster than how they are normally performed. The previous literature confirmed that the specific behavioural cues we observed are used when expressing anger (Ekman et al., 1972; Ekman, 1992). Moreover, re-search on body language suggests that specific hands and arms gestures that we observed such as pointing and arms swinging displayed during a conversation are also reliable indicators of anger (Saha et al., 2014; Ajibo et al., 2020), thus supporting our interpretation.

We grouped into Category 3 the behaviours that indicate a state of sufferance, physical pain and the presence of injuries caused by the aggressive encounter. They refer to the postures that a person assumed in order to sustain his/her body (“Leaning forced”, “Sitting forced”; Figure 1) and the self-caring activities performed to medicate visible injuries (“Self-care”; Figure 1). Since specific facial expressions and movements allowed us to recognize when the antagonists were crying, we also included this state into Category 3. All the behaviours we listed in this category have been described as reliable non-verbal indicators of pain and distress in previous literature (von Baeyer et al., 1984; Hadjistavropoulos & Craig, 2002).

The behaviours we observed and interpreted as expression of anxiety, anger, and physical sufferance find some correspondence with the reactions reported in nonhuman primates species in the post-conflict context. In the aftermath of a conflict, non-human primates express a state of anxiety by increasing their rate of self-scratching and self-grooming (Baker & Aureli, 1997; Fraser et al., 2008; Palagi et al., 2014). The occurrence of such self-directed behaviours is due to the hormonal response elicited by the stressful stimulus, i.e., the conflict, which increases the production of glucocorticoids inducing in turn the occurrence of scratching as a behavioural response (Schino et al., 1996; Troisi, 2002; Sanders & Akiyama 2018; Rinaldi, 2019; Sapolsky, 2021). Similarly, the hormonal circuit activated in humans as a reaction to a stressful stimulus generates a behavioural response the consists in the increased frequency of occurrences of displacement activities (Troisi, 2002; Steimer, 2022) and also modulates the predisposition to behave aggressively, thus explaining the increase in aggressive actions as a consequence of social stress (Veenema & Neumann, 2007; Neumann et al., 2010). The expression of anger after the experience of being involved in a conflict might represent for the previous antagonist a form of redirected aggression, which in human and nonhuman primates has been interpreted as a relief for emotional arousal as it offers an outlet to antagonists’ frustration (Fitz, 1976; Marcus-Newhall, 2000; Kazem & Aureli, 2005; McKittrick et al., 2009; Romero et al., 2011; Pallante et al., 2018).

Differently from the first three categories, the behavioural patterns included in Category 4 have been frequently observed in the post-conflict context but found no correspondence with any specific emotional state. They refer to specific postures (“Hand on hips”, “Hand in pockets”, “Arm crossed”, “Leaning”; Figure 3) and interactions (“Talking with gestures”; Figure 3). Since no studies previously linked the behaviours, we included in Category 4 with the expression of distress, we avoided any interpretation and labelled this category “Unclassified behaviours”. However, body postures and hands gestures displayed during a conversation—as we described in the case of “Talking with gestures”—may be related to specific emotional states and convey a precise meaning in terms of non-verbal communication (Ekman, 1965; Shan et al., 2007; de Gelder, 2009; Baltrušaitis et al., 2011; Hostetter, 2011). In order to evaluate if the behaviours of Category 4 are typical of a context of distress, we suggest that future research introduce a comparison with a baseline condition. In nonhuman primates research, an observational procedure consisting of a controlled design has been developed to assess the link between the expression of a specific behaviour with a specific context (de Waal & Yoshihara, 1983; Fraser et al., 2008). This method allowed us to understand that some interactions or behavioural patterns that an individual displays in several different contexts increase their expression in a specific situation. For instance, in order to evaluate whether antagonists of a conflict experience a state of distress in its aftermath, primatologists compared the frequency (e.g., for events, such as a self-scratching) and the duration (e.g., for a state, such as a self-grooming) of self-directed behaviours, whose occurrence is not limited to the post-conflict context, in presence and absence of conflicts. Such comparison revealed that an increase in the expression of self-directed behaviours characterizes the post-conflict context, suggesting therefore an emotional variation (Baker & Aureli, 1997; Fraser et al., 2008; Palagi et al., 2014). Similarly, comparing the occurrence of the behaviours included in Category 4 with a baseline context which is not characterized by any conflict would reveal whether their expression is typical of the post-conflict context, underlying their link with a state of distress.

The second aim of the study was to use the ethogram to quantify the distress-related behaviours in real-life events in our sample of CCTV footage. This quantitative coding allowed us to detect which behaviours where more frequently displayed and which occurred more rarely. The results obtained from the correlation matrix revealed that except for “Sitting forced” and “Leaning forced”, there are no specific behaviours expressed in the aftermath of a conflict displayed in combination with each-other. Along the same lines, a PCA revealed a co-occurrence of behaviours that does not result in any clear cluster, since the percentage of the variance explained by each dimension obtained is too low to convey any meaningful information. An interesting though weak association was observed between some of the behaviours of Category 4 and Category 1 (e.g., “Armed crossed” and “Pacing” or “Moving on the place” and “Talking with gestures”). This relationship suggests that some behaviours that have not been previously identified as expression of distress might be associated to a state of anxiety. However, the absence of any significant co-occurrence of behaviours resulting from the PCA highlights that antagonists in the post-conflict context express in general a state of distress by displaying a combination of behaviours that is not fixed. Several of the behaviours we observed might therefore have a similar function and they can replace each other to express distress (e.g., both Self-touching and Redirected attention can be displayed to express anxiety). By conveying a similar message, these behaviours might thus function as substitutes, as it occurs with the redundancy of signals characterizing communicative systems (Krebs & Dawkins, 1984). Our results reveal that these behaviours do not appear together, suggesting a variability in how people express distress. However, future analysis shall be run to confirm our hypothesis by testing the co-occurrence of the behaviours that express the same emotional state following a theoretical driven approach on the basis of the qualitative categorization we provided in our study.

Compared to the methods traditionally employed in the study of distress that were limited to experimental settings, the use of CCTV footage allowed us to observe naturally occurring events characterized by tension. However, we can only assume from the context that the antagonists that we took into consideration actually experienced a feeling of distress, and therefore the relation between the behaviours we documented, and the underpinning emotional state remains here only speculative. This is similar to the assumption that has been made in the previous research (Barash, 1974; Shreve et al., 1988). Further measurements on the behaviours in the ethogram in controlled settings could reinforce our conclusions both by providing such physiological association and by investigating their presence in different contexts of tension.

The ethological approach to the study of distress that we adopted further allows to collect and process data that are suitable for cross-species comparisons, introducing the possibility to formulate biological explanations on human behaviour, whose study is limited in the social sciences. Although a shared research method does not necessarily imply the observation of the same phenomena, it enables different disciplines to disclose the existence of events, behaviours and patterns typically investigated in different fields that would not be evident otherwise (Martin, 2017; Nippert-Eng, 2015). Until now, a shared approach has seldom been considered both in the fields of social sciences, where evolutionary explanations are frequently neglected, and in the ethological research, whose focus is usually skewed towards nonhuman species (Ellis, 1996; Massey, 2002). As a consequence, this separation has led to the formulation of different explanations for the same phenomena investigated in the two fields (Kavish et al., 2017). By combining methods of different disciplines, our work aims to be a further step in the development of a more inclusive perspective.

The data we analysed are purely behavioural and they were restricted to visual information, since the sound was excluded from the footage. The recording of the voice would indeed have reinforced our interpretation of specific social interactions as related to distress, such as talking with gestures or talking with aggressive gestures. Similarly, we were not able to detect distress expressed through vocalizations or verbal cues only. Therefore, given the role of vocal indicators in the expression of emotions (Hall & Knapp, 2013; Bänziger et al., 2014; Scherer et al., 2011), we encourage future research to consider the potential effect that distress-related sounds could play on the evaluation of tension in the post-conflict context.

Limitations of our study are linked to the period of data collection and the location of the cameras. All the conflict events we analysed were recorded during the COVID-19 pandemic, when the Dutch government advised for specific behaviours to be adopted at individual (i.e., improving hand hygiene and limiting the touch of specific parts of the body, such as the face, that could increase the risk of contracting the infection) and relational level (i.e., keeping 1. 5 m distance). Such measures may therefore have constrained people’s behaviour, limiting their propensity to self-touch, and their motivation to socially interact, as for talking with gestures. In terms of space, the location of the camera in the city of Amsterdam restricts our sample to a single city and calls into questions the generalizability of our conclusions. Although the universality in the expression of basic emotions has been accepted by the scientific community, a cultural variation may still persist in relation to specific gestures displayed to communicate emotions (Darwin, 2009 [1872]; Ekman & Friesen, 1971; Kret et al., 2022). Therefore, we cannot claim to what extent the behaviours we identified as indicators of distress may represent socially learned conventions instead of inherited predispositions, and as a consequence if our limited sample may in turn be generalizable to other cultural contexts.

Our first step in the detection of the behavioural correlates of distress opens possibilities for future investigations on the effects of distress in group’s social dynamics in the post-conflict context and to use the ethogram to implement conflict mitigation strategies. Recent studies documented that human bystanders play an active role in managing conflicts in public (Philpot et al., 2020). However, the factors driving bystander interventions are still largely unexplored and we do not currently know whether, besides the relational and context characteristics that trigger the intervention (Liebst et al., 2019, Lindegaard et al., 2021), the behavioural cues indicating the emotional state of the antagonists are also involved. Signs of distress may communicate the emotional state of the antagonists to bystanders. Therefore, the ethogram we provided can be used in research on interpersonal conflicts to understand whether bystanders behaviour is actually affected by the observation of the behavioural cues of distress of the antagonists, and if such awareness facilitates in turn bystanders’ response and subsequent mitigation interventions. Similarly, in nonhuman primate species the expression of distress in the post-conflict context leads to bystander intervention, which has a calming effect in the receiver (Clay et al., 2018).

A further and more operative application of the ethogram consists of informing people involved or attending an incident on which behaviours are linked to a state of distress, indicating how to read (in the case of bystanders) and communicate (in the case of antagonists) distress. On the one hand, acknowledging what the behavioural signs of distress are helps bystanders to elaborate an appropriate response to a person in need of help. On the other hand, and from the antagonist’s perspective, a clear expression of distress might convey a potential need for help, hence actively soliciting bystanders’ interventions. The informative value of nonverbal behavioural indicators of distress have proven to be effective in humans in contexts such as patient-medical consultations (Shreve et al., 1988; Zimmermann et al., 2011) and long-term risk detection from ambulance workers (Halpern et al., 2011). For instance, experimental research reported that the expression of fear and anger elicited, respectively, affiliation and avoidance in the observers, suggesting that their reaction is modulated according to the perceived emotion (Marsh et al., 2005). Therefore, the ethogram we developed can be used to implement the recognition of the behaviours indicating distress, which may help non-professional third parties, such as guardians, and professionals such as police and first aid personal, in providing the appropriate response. The application of the ethogram to recognize the cues of distress is not limited to a human user. Computer vision systems are increasingly implemented to detect human behaviours; therefore, a further application might consist in developing algorithms to recognize the cues of distress in order to identify emergency situations through surveillance cameras in public spaces (Bernasco et al., 2021). Such automated coding procedure has the potential to run a large-scale analysis of the video content which until now has been constrained by manual coding. With the tool proposed in the present study, we hope to facilitate the understanding of the social dynamics in a context of violence and, more importantly, its management strategies.

The observation of human emotional behaviour has been traditionally confined to experimental settings, frequently becoming uncoupled from real life situations. With the intent to provide an ecologically valid picture of human behavioural repertoire as a response to a distressful event, we draw from the ethological method to analyse video of interpersonal conflicts of human adults occurring in public spaces. We developed the ethogram of the behaviours observed, tested it for the inter-coder reliability, and used it to explore the behavioural variability expressed in the post-conflict context. We observed that in humans a conflict event elicits the expression of behaviours indicating a state of distress, in particular anxiety, anger, and physical sufferance. These findings are consistent with previous research that documented a similar response in other nonhuman primate species, thus highlighting a phylogenetic continuity. We encourage the use of ethogram for the systematic study of human behaviour in real life situations and to understand the expression of distress in a comparative perspective.

## ACKNOWLEDGMENTS

We wish to thank Lucia Berti for the drawings, Hans Myhre Sunde and Wim Bernasco for helping with the inter-coder reliability, and Otto Adang for valuable discussion on the manuscript. This study was funded by the Dutch Research Council (NWO VI.Vidi.195.083) by means of a grant awarded to Marie Rosenkrantz Lindegaard. This project has also received funding from the European Union’s Horizon 2020 research and innovation program under the Marie Sklodowska-Curie grant agreement No. 101029161 awarded to Virginia Pallante.

## AUTHOR CONTRIBUTIONS

Conceptualization: V.P. and M.R.L. Methodology: V.P., P.E.-E., and M.R.L. Formal Analysis: V.P. and P.E.-E. Investigation: V.P., P.E.-E., and M.R.L. Data Curation: V.P., P.E.-E., and M.R.L. Writing—Original Draft Preparation: V.P., P.E.-E., and M.R.L. Writing—Review and Editing: V.P., P.E.-E., and M.R.L. Supervision: M.R.L. Project Administration: M.R.L. Funding Acquisition: V.P. and M.R.L. All authors have read and agreed to the published version of the manuscript.

## REFERENCES

1. Ajibo, C.A., Ishi, C.T., Mikata, R., Liu, C. & Ishiguro, H. (2020). Analysis of body gestures in anger expression and evaluation in android robot. – Adv. Robot. 34: 1581–1590. https://doi.org/10.1080/01691864.2020.1855244

2. Altmann, J. (1974). Observational study of behavior: sampling methods. – Behaviour 49: 227–266. doi:https://doi.org/10.1163/156853974X00534

3. Argyle, M. (2013). Bodily communication. – Routledge.

4. Arsenio, W.F. & Killen, M. (1996). Conflict-related emotions during peer disputes. Early Educ. Devel. – 7: 43–57. https://doi.org/10.1207/s15566935eed0701_4

5. Aureli, F. & Schaik, C.P.V. (1991). Post-conflict behaviour in long-tailed macaques (Macaca fascicularis) II. Coping with the uncertainty. – Ethology 89: 101–114. https://doi.org/10.1111/j.1439-0310.1991.tb00297.x

6. Aureli, F. & de Waal, F.B.M., eds (2000). Natural Conflict Resolution. – University of California Press, Berkeley, CA, USA.

7. Baker, K.C. & Aureli, F. (1997). Behavioural indicators of anxiety: An empirical test in chimpanzees. – Behaviour 134: 1031–1050. https://www.jstor.org/stable/4535489

8. Baltrušaitis, T., McDuff, D., Banda, N., Mahmoud, M., El Kaliouby, R., Robinson, P. & Picard, R. (2011). Real-time inference of mental states from facial expressions and upper body gestures. – In: 2011 IEEE International Conference on Automatic Face & Gesture Recognition (FG). 21–23 March 2011. IEEE, Santa Barbara, CA, USA, p. 909-914. DOI:10.1109/FG.2011.5771372

9. Bänziger, T., Patel, S. & Scherer, K.R. (2014). The role of perceived voice and speech characteristics in vocal emotion communication. – J. Nonverbal Behav. 38: 31–52. https://doi.org/10.1007/s10919-013-0165-x

10. Barash, D.P. (1974). Human ethology: Displacement activities in a dental office. – Psychol. Rep. 34: 947–949. https://doi.org/10.2466/pr0.1974.34.3.947

11. Bardi, M., Shimizu, K., Barrett, G. M., Borgognini-Tarli, S.M. & Huffman, M.A. (2003). Peripartum cortisol levels and mother–infant interactions in Japanese macaques. – Am. J. Phy. Anthropol. 120: 298–304. https://doi.org/10.1002/ajpa.10150

12. Bardi, M., French, J.A., Ramirez, S.M. & Brent, L. (2004). The role of the endocrine system in baboon maternal behavior. – Biol. Psych. 55: 724–732. https://doi.org/10.1016/j.biopsych.2004.01.002

13. Bardi, M., Koone, T., Mewaldt, S. & O’Connor, K. (2011). Behavioral and physiological correlates of stress related to examination performance in college chemistry students. – Stress 14: 557–566. https://doi.org/10.3109/10253890.2011.571322

14. Bernasco, W., Hoeben, E.M., Koelma, D., Liebst, L.S., Thomas, J., Appelman, J., Snoek, C.G.M. & Lindegaard, M.R. (2021). Promise into practice: Application of computer vision in empirical research on social distancing. – Sociol. Methods Res. https://doi.org/10.1177/00491241221099554.

15. Butovskaya, M. & Kozintsev, A. (1999). Aggression, friendship, and reconciliation in Russian primary schoolchildren. – Aggress. Behav. 25: 125–139. https://doi.org/10.1002/(SICI)1098-2337(1999)25:2<125::AID-AB5>3.0.CO;2-X

16. Castles, D.L. & Whiten, A. (1998). Post-conflict behaviour of wild olive baboons. II. Stress and self-directed behaviour. – Ethology 104: 148–160. https://doi.org/10.1111/j.1439-0310.1998.tb00058.x

17. Clay, Z. & de Waal, F.B.M. (2013). Bonobos respond to distress in others: Consolation across the age spectrum. – PLoS ONE 8: e55206.

18. Clay, Z., Palagi, E. & de Waal, F. B. M. (2018). Ethological approaches to empathy in primates. – In: Neuronal correlates of empathy. Academic Press, p. 53-66. https://doi.org/10.1016/B978-0-12-805397-3.00005-X

19. Darwin, C. (2009 [1872]). The Expression of the Emotions in Man and Animals. – HarperCollinsPublishers.

20. Das, M., Penke, Z. & Van Hooff, J.A. (1998). Postconflict affiliation and stress-related behavior of long-tailed macaque aggressors. – Int. J. Primatol. 19: 53–71. https://doi.org/10.1023/A:1020354826422

21. de Gelder, B. (2009). Why bodies? Twelve reasons for including bodily expressions in affective neuroscience. – Phil. Trans. Royal Soc. B Biol. Sci. 364: 3475–3484. https://doi.org/10.1098/rstb.2009.0190

22. de Waal, F.B.M. & Yoshihara, D. (1983). Reconciliation and redirected affection in rhesus monkeys. – Behaviour 85: 224–241. https://doi.org/10.1163/156853983X00237

23. Eibl-Eibesfeldt, I. (2007). Human Ethology. – Aldine Transaction, New Brunswick, N.J.

24. Ejbye-Ernst, P., Lindegaard, M.R. & Bernasco, W. (2020). A CCTV-based analysis of target selection by guardians intervening in interpersonal conflicts. – Europ. J. Criminol. 19: 1260–1279. https://doi.org/10.1177/1477370820960338

25. Ekman, P. (1965). Differential communication of affect by head and body cues. – J. Person. Soc. Psychol. 2: 926–735. doi:10.1037/h0022736

26. Ekman P. (1992). Are there basic emotions? – Psychol. Rev. 99: 5509–553. https://doi.org/10.1037/0033-295X.99.3.550

27. Ekman, P. (1998). Universality of emotional expression? A personal history of the dispute. – In: Darwin, C.(2009 [1872]). The Expression of the Emotions in Man and Animals, 3rd edition. HarperCollinsPublishers, p. 363-393.

28. Ekman, P. & Friesen, W.V. (1971). Constants across cultures in the face and emotion. – J. Pers. Soc. Psychol. 17: 124–129. https://doi.org/10.1037/h0030377

29. Ekman, P., Friesen, W.V. & Ellsworth, P. (1972). Emotion in the Human Face: Guide-lines for Research and an Integration of Findings. – Pergamon Press, Pergamon, New York.

30. Ellis, L., (1996). A discipline in peril: Sociology’s future hinges on curing its biophobia. – Am. Sociol. 27: 21–41. https://doi.org/10.1007/BF02692016

31. Fitz, D. (1976). A renewed look at Miller’s conflict theory of aggression displacement. – J. Pers. Soc. Psychol. 33: 725. https://doi.org/10.1037/0022-3514.33.6.725

32. Fraser, O.N., Stahl, D. & Aureli, F. (2008). Stress reduction through consolation in chimpanzees. – PNAS 105: 8557–8562. https://doi.org/10.1073/pnas.080414110.

33. Fujisawa, K.K., Kutsukake, N. & Hasegawa, T. (2005). Reconciliation pattern after aggression among Japanese preschool children. – Aggress. Behav. 31: 138–152. https://doi.org/10.1002/ab.20076

34. Hadjistavropoulos, T. & Craig, K.D. (2002). A theoretical framework for understanding self-report and observational measures of pain: a communications model. – Behav. Res. Ther. 40: 551–570. https://doi.org/10.1016/S0005-7967(01)00072-9

35. Hall, J. A. & Knapp, M. L. (2013). Nonverbal communication. –de Gruyter Mouton, Boston.

36. Halpern, J., Maunder, R. G., Schwartz, B. & Gurevich, M. (2011). Identifying risk of emotional sequelae after critical incidents. – Emerg. Med. J. 28: 51–56. http://dx.doi.org/10.1136/emj.2009.082982

37. Heesen, R., Austry, D.A. & Upton, Z.; Clay, Z. (2022). Flexible signalling strategies by victims mediate post-conflict interactions in bonobos. – Philos. T. R. Soc. B 377: 20210310. https://doi.org/10.1098/rstb.2021.0310.

38. Hostetter, A.B. (2011). When do gestures communicate? A meta-analysis. – Psychol. Bull. 137: 297–315. https://doi.org/10.1037/a0022128

39. Jones, L.K., Jennings, B.M., Goelz, R.M., Haythorn, K.W., Zivot, J.B. & de Waal, F.B.M. (2016). An ethogram to quantify operating room behavior. – Ann. Behav. Med. 50: 487–496. https://doi.org/10.1007/s12160-016-9773-0

40. Kalueff, A.V., La Porte, J.L. & Bergner, C.L., (2010). Neurobiology of grooming behaviour. – Cambridge University Press, Cambridge, MA.

41. Kavish, N., Fowler-Finn, K. & Boutwell, B.B. (2017). Criminology’s modern synthesis: remaking the science of crime with Darwinian insight. – In: The evolution of psychopathology. Springer, Cham, p. 171–183.

42. Kazem, A.J.N. & Aureli, F. (2005). Redirection of aggression: Multiparty signalling within a network? – In: Animal Communication Net-works (McGregor, P.K., ed.). Cambridge University Press, Cambridge, UK, p. 191–218.

43. Klusemann, S. (2009). Atrocities and confrontational tension. – Front. Behav. Neurosci. 3: 42. https://doi.org/10.3389/neuro.08.042.2009

44. Kret, M.E., Massen, J.J. & de Waal, F. (2022). My fear is not, and never will be, your fear: On emotions and feelings in animals. – Affect. Sci. 3: 182–189. https://doi.org/10.1007/s42761-021-00099-x

45. Kutsukake, N. & Castles, D.L. (2001). Reconciliation and variation in post-conflict stress in Japanese macaques (Macaca fuscata fuscata): testing the integrated hypothesis. – Anim. Cogn., 4: 259–268. https://doi.org/10.1007/s10071-001-0119-2

46. Landis, J.R. & Koch, G.G. (1977). The measurement of observer agreement for categorical data. – Biometrics. 33: 159–174. https://doi.org/10.2307%2F2529310

47. Leavens D, Aureli F, Hopkins W. (2004). Behavioral evidence for the cutaneous expression of emotion in a chimpanzee (Pan Troglodytes). – Behaviour 141: 979–997. doi: https://doi.org/10.1163/1568539042360189

48. Lehner, P.N. (1998). Handbook of ethological methods. – Cambridge University Press.

49. Levine, M., Taylor, P.J. & Best, R. (2011). Third parties, violence, and conflict resolution: The role of group size and collective action in the microregulation of violence. – Psychol. Sci. 22: 406–412. https://doi.org/10.1177/0956797611398495

50. Liebst, L.S., Philpot, R., Bernasco, W., Dausel, K.L., Ejbye-Ernst, P., Nicolaisen, M.H. & Lindegaard, M.R. (2019). Social relations and presence of others predict bystander intervention: Evidence from violent incidents captured on CCTV. – Aggress. Behav. 45: 598–609. https://doi.org/10.1002/ab.21853

51. Lindegaard, M.R., Liebst, L.S., Bernasco, W., Heinskou, M.B., Philpot, R., Levine, M. & Verbeek, P. (2017). Consolation in the aftermath of robberies resembles post-aggression consolation in chimpanzees. – PloS one 12: e0177725. https://doi.org/10.1371/journal.pone.0177725

52. Lindegaard, M. R. & Bernasco, W. (2018). Lessons learned from crime caught on camera. – J. Res. Crime Delinq. 55: 155–186. https://doi.org/10.1177/0022427817727830

53. Lindegaard, M.R., Liebst, L. S., Philpot, R., Levine, M. & Bernasco, W. (2021). Does Danger Level Affect Bystander Intervention in Real-Life Conflicts? Evidence From CCTV Footage. – Soc. Psychol. Pers. Sci. 13: 795–802. https://doi.org/10.1177/19485506211042683

54. Ljungberg, T., Westlund, K. & Forsberg, A.J.L. (1999). Conflict resolution in 5-year-old boys: does postconflict affiliative behaviour have a reconciliatory role?. – Anim. Behav. 58: 1007–1016. https://doi.org/10.1006/anbe.1999.1236

55. Maestripieri, D., Schino, G., Aureli, F. & Troisi, A. (1992). A modest proposal: displacement activities as an indicator of emotions in primates. – Anim. Behav. 44: 967–979. https://doi.org/10.1016/S0003-3472(05)80592-5

56. Marcus-Newhall, A., Pedersen, W. C., Carlson, M. & Miller, N. (2000). Displaced aggression is alive and well: a meta-analytic review. – J. Pers. Soc. Psychol. 78: 670. https://doi.org/10.1037/0022-3514.78.4.670

57. Marsh, A. A., Ambady, N. & Kleck, R. E. (2005). The effects of fear and anger facial expressions on approach-and avoidance-related behaviors. – Emotion 5: 119. https://doi.org/10.1037/1528-3542.5.1.119

58. Martin, J.L. (2017). Thinking through methods: A social science primer. – University of Chicago Press.

59. Massey, D.S. (2002). A brief history of human society: The origin and role of emotion in social life. – Am. Social. Rev. 67: 1–29. https://doi.org/10.2307/3088931

60. McKittrick, C.R., Blanchard, D.C., Hardy, M.P. & Blanchard, R.J. (2009). Social stress effects on hormones, brain, and behaviour. – In: Hormones, brain and behaviour (Pfaff, D.W., Arnold, A.P, Etgen, A.M., Fahrbach, S.E. & Rubin, R.T., eds.). Academic, San Diego, CA, p. 341–347. doi:10.1016/B978-008088783-8.00009-7.

61. Mohiyeddini, C. & Semple, S. (2013). Displacement behaviour regulates the experience of stress in men. – Stress 16: 163–171. https://doi.org/10.3109/10253890.2012.707709

62. Mosselman, F., Weenink, D. & Lindegaard, M.R. (2018). Weapons, body postures, and the quest for dominance in robberies: A qualitative analysis of video footage. – J. Res. Crime Delinq. 55: 3–26. https://doi.org/10.1177/0022427817706525

63. Nassauer, A. (2016). From peaceful marches to violent clashes: A micro-situational analysis. – Soc. Mov. Stud., 15: 515–530. https://doi.org/10.1080/14742837.2016.1150161

64. Nassauer, A. (2018). How robberies succeed or fail: Analyzing crime caught on CCTV. – J. Res. Crime Delinq, 55: 125–154. https://doi.org/10.1177/0022427817715754

65. Neumann, I.D., Veenema, A.H. & Beiderbeck, D. (2010). Aggression and anxiety: Social context and neurobiological links. – Front. Behav. Neurosci. 12. https://doi.org/10.3389/fnbeh.2010.00012.

66. Niaura, R., Shadel, W.G., Britt, D.M. & Abrams, D.B. (2002). Response to social stress, urge to smoke, and smoking cessation. – Addict. Behav. 27: 241–250. https://doi.org/10.1016/S0306-4603(00)00180-5

67. Nippert-Eng, C. (2015). Watching closely: A guide to ethnographic observation. – Oxford University Press.

68. Palagi, E., Dall’Olio, S., Demuru, E. & Stanyon, R. (2014). Exploring the evolutionary foundations of empathy: consolation in monkeys. – Evol. Hum. Behav. 35: 341–349. https://doi.org/10.1016/j.evolhumbehav.2014.04.002

69. Pallante, V., Stanyon, R. & Palagi, E. (2018). Calming an aggressor through spontaneous post-conflict triadic contacts: Appeasement in Macaca tonkeana. – Aggress. Behav. 44: 406–415. https://doi.org/10.1002/ab.21761

70. Philpot, R., Liebst, L.S., Møller, K.K., Lindegaard, M.R. & Levine, M. (2019). Capturing violence in the night-time economy: A review of established and emerging methodologies. – Aggress. Violent Behav. 46: 56–65. https://doi.org/10.1016/j.avb.2019.02.004

71. Philpot, R., Liebst, L.S., Levine, M., Bernasco, W. & Lindegaard, M.R. (2020). Would I be helped? Cross-national CCTV footage shows that intervention is the norm in public conflicts. – Am. Psychol. 75: 66. https://doi.org/10.1037/amp0000469

72. Philpot, R., Liebst, L.S., Lindegaard, M.R., Verbeek, P. & Levine, M. (2022). Reconciliation in human adults: A video-assisted naturalistic observational study of post conflict conciliatory behaviour in interpersonal aggression. – Behaviour 159: 1225–1261.

73. Pico-Alfonso, M.A., Mastorci, F., Ceresini, G., Ceda, G.P., Manghi, M., Pino, O. & Sgoifo, A. (2007). Acute psychosocial challenge and cardiac autonomic response in women: The role of estrogens, corticosteroids, and behavioral coping styles. – Psychoneuroendocrinology 32: 451-463. https://doi.org/10.1016/j.psyneuen.2007.02.009

74. Rinaldi, G. (2019). The itch-scratch cycle: A review of the mechanisms. – Dermatol. Pract. Concep. 9: 90–97. https://doi.org/10.5826/dpc.0902a03.

75. Romero, T., Castellanos, M.A. & de Waal, F.B.M. (2011). Post-conflict affiliation by chimpanzees with aggressors: Other-oriented versus selfish political strategy. – PLoS ONE 6: e22173. https://doi.org/10.1371/journal.pone.0022173

76. Saha, S., Datta, S., Konar, A. & Janarthanan, R. (2014). A study on emotion recognition from body gestures using Kinect sensor. – In: 2014 international conference on communication and signal processing. IEEE, p. 056–060. DOI:10.1109/ICCSP.2014.6949798

77. Sanders, K.M. & Akiyama, T. (2018). The vicious cycle of itch and anxiety. – Neurosci. Biobehav. Rev. 87: 17–26. https://doi.org/10.1016/j.neubiorev.2018.01.009.

78. Sapolsky, R.M. (2021). Glucocorticoids, the evolution of the stress-response, and the primate predicament. – Neurobiol. Stress 14: 100320. https://doi.org/10.1016/j.ynstr.2021.100320

79. Scherer, K.R., Clark-Polner, E. & Mortillaro, M. (2011). In the eye of the beholder? Universality and cultural specificity in the expression and perception of emotion. – Int. J. Psychol. 46: 401–435. https://doi.org/10.1080/00207594.2011.626049

80. Schino, G., Perretta, G., Taglioni, A. M., Monaco, V. & Troisi, A. (1996). Primate displacement activities as an ethopharmacological model of anxiety. – Anxiety, 2: 186–191. https://doi.org/10.1002/(SICI)1522-7154(1996)2:4<186::AID-ANXI5>3.0.CO;2-M

81. Sgoifo, A., Braglia, F., Costoli, T., Musso, E., Meerlo, P., Ceresini, G. & Troisi, A. (2003). Cardiac autonomic reactivity and salivary cortisol in men and women exposed to social stressors: relationship with individual ethological profile. – Neurosci. Biobehav. Rev. 27: 179–188. https://doi.org/10.1016/S0149-7634(03)00019-8

82. Shan, C., Gong, S. & McOwan, P.W. (2007). Beyond Facial Expressions: Learning Human Emotion from Body Gestures. – In: BMVC. p. 1–10. doi:10.5244/C.21.43

83. Shreve, E.G., Harrigan, J.A., Kues, J.R., & Kagas, D.K. (1988). Nonverbal expressions of anxiety in physician-patient interactions. – Psychiatry 51: 378–384. https://doi.org/10.1080/00332747.1988.11024414

84. Steimer, T. (2022). The biology of fear-and anxiety-related behaviors. – Dialog. Clin. Neurosci. 4: 231–249. https://doi.org/10.31887/DCNS.2002.4.3/tsteimer.

85. Troisi, A. (1999). Ethological research in clinical psychiatry: the study of nonverbal behavior during interviews. – Neurosci. Biobehav. Rev. 23: 905–913. https://doi.org/10.1016/S0149-7634(99)00024-X

86. Troisi, A. (2002). Displacement activities as a behavioral measure of stress in nonhuman primates and human subjects. – Stress 5: 47–54. https://doi.org/10.1080/102538902900012378

87. Troisi, A., Delle Chiaie, R., Russo, F., Russo, M.A., Mosco, C. & Pasini, A. (1996). Nonverbal behavior and alexithymic traits in normal subjects: individual differences in encoding emotions. – J. Nerv. Ment. Dis. 184: 561–566. https://doi.org/10.1097/00005053-199609000-00008

88. Veenema, A.H. & Neumann, I.D. (2007). Neurobiological mechanisms of aggression and stress coping: A comparative study in mouse and rat selection lines. – Brain Behav. Evolut. 70: 274–285. https://doi.org/10.1159/000105491.

89. von Baeyer, C.L., Johnson, M.E. & McMillan, M.J. (1984). Consequences of nonverbal expression of pain: Patient distress and observer concern. – Soc. Sci. Med. 19: 1319–1324. https://doi.org/10.1016/0277-9536(84)90019-4

90. Westlund, K., Horowitz, L., Jansson, L. & Ljungberg, T. (2008). Age effects and gender differences on post-conflict reconciliation in preschool children. – Behaviour 145: 1525–1556. https://doi.org/10.1163/156853908786131351

91. Whitehouse, J., Milward, S.J., Parker, M.O., Kavanagh, E. & Waller, B.M. (2022). Signal value of stress behaviour. – Evol. Hum. Behav. 43: 325–333. https://doi.org/10.1016/j.evolhumbehav.2022.04.001

92. Zandara, M., Villada, C., Hidalgo, V. & Salvador, A. (2018). Assessing the antecedents and consequences of threat appraisal of an acute psychosocial stressor: the role of optimism, displacement behavior, and physiological responses. – Stress 21: 304–311. https://doi.org/10.1080/10253890.2018.1449830

93. Zimmermann, C., Del Piccolo, L., Bensing, J., Bergvik, S., De Haes, H., Eide, H. & Finset, A. (2011). Coding patient emotional cues and concerns in medical consultations: the Verona coding definitions of emotional sequences (VR-CoDES). – Patient Educ. Couns. 82: 141–148. https://doi.org/10.1016/j.pec.2010.03.017

