## Supplementary figures and images for "An ethogram method for the analysis of human distress in the aftermath of public conflicts"

### Supplementary Figure 1

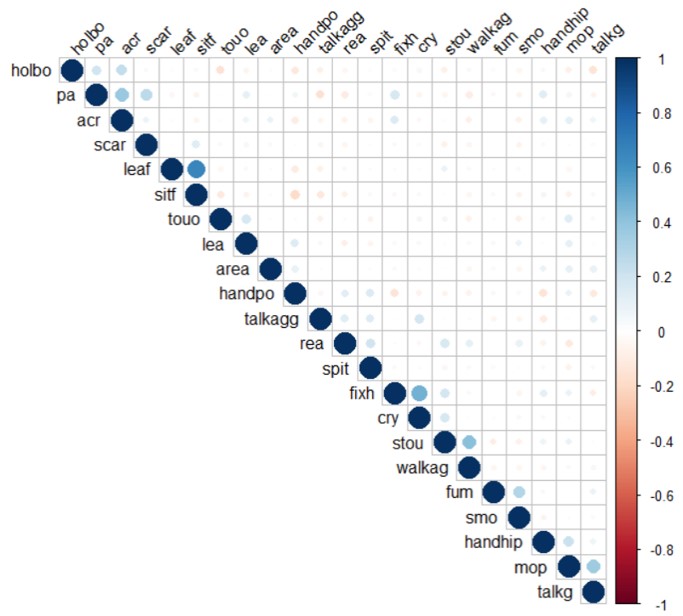

### Supplementary Figure 2

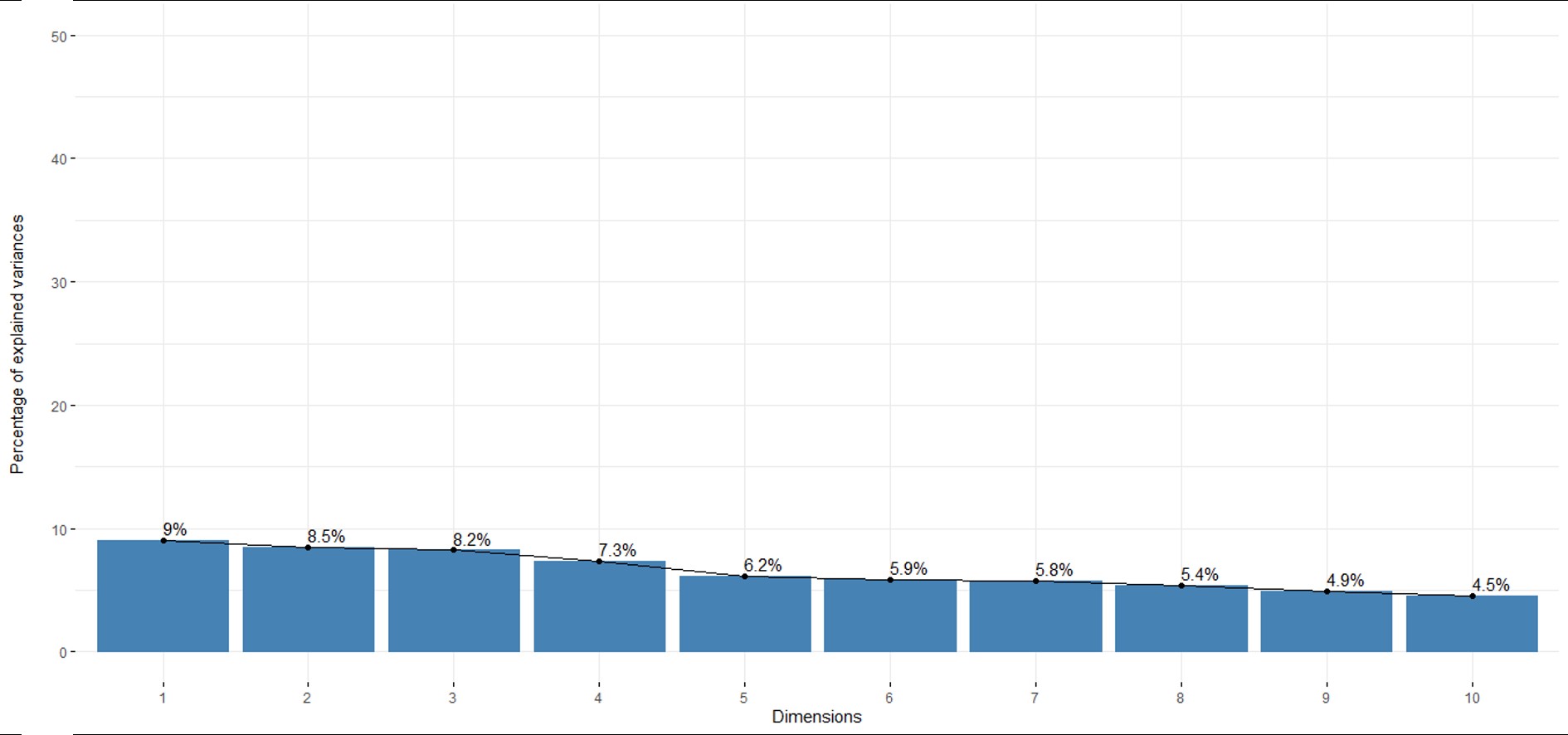

### Supplementary Figure 3

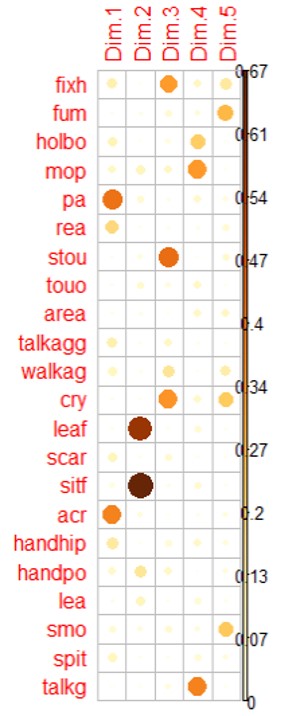

### Supplementary Figure 4

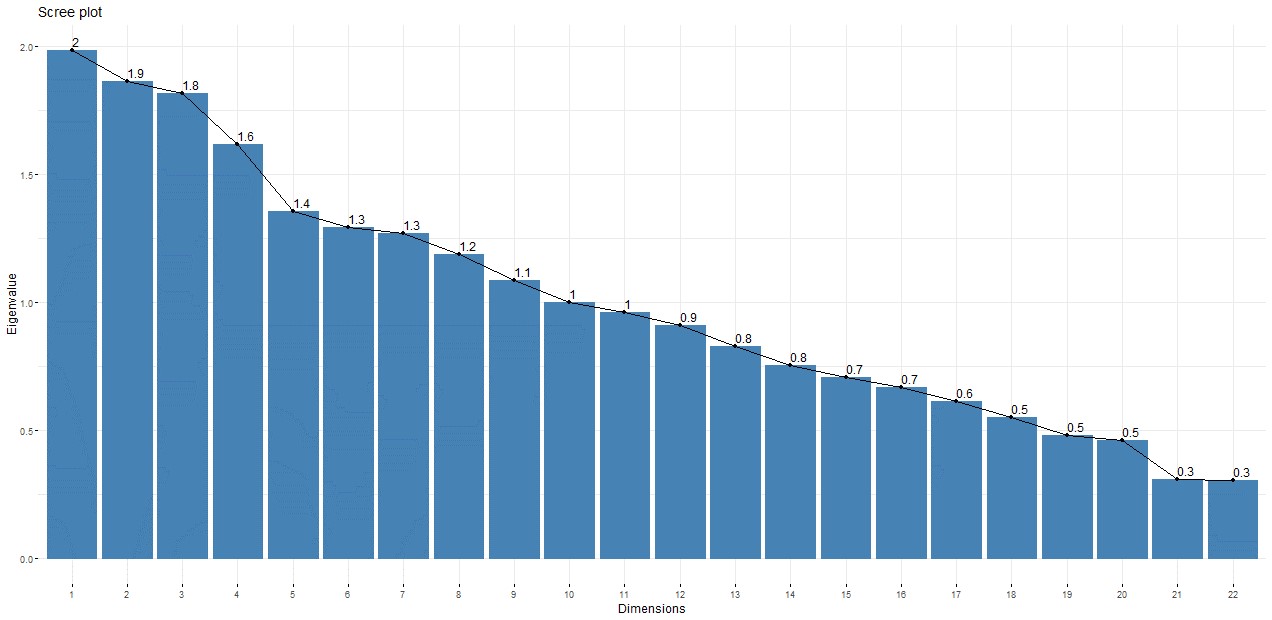
