## Supplementary Table 1 for "An ethogram method for the analysis of human distress in the aftermath of public conflicts"

**Table S1.** Cohen's Kappa (κ) for each behavioural code.

| **Behaviour** | **Cohen's Kappa (κ)** |
|---|---|
| acr | 1 |
| area | 1 |
| cry | 0.655 |
| fixh | 0.660 |
| fum | 0.660 |
| handpo | 0.817 |
| handhip | 1 |
| holbo | 1 |
| lea | 0.736 |
| leaf | 0.901 |
| mop | 0.610 |
| pa | 0.745 |
| rea | 0.642 |
| scar | 0.655 |
| stou | 0.722 |
| sitf | 0.685 |
| smo | 0.784 |
| spi | 0.647 |
| talkg | 0.790 |
| touo | 0.620 |
| talkagg | 0.697 |
| walkag | 0.655 |
