## Supplementary Table 2 for "An ethogram method for the analysis of human distress in the aftermath of public conflicts"

**Table S2**. Correlation matrix analysis. Different colours are indicative of the category to which each behaviour belongs.

| | fixh | fum | holbo | mop | pa | rea | smo | stou | touo | area | talkagg | walkag | cry | leaf | scar | sitf | acr | handhip | handpo | lea | spit | talkg |
|---|---|---|---|---|---|---|---|---|---|---|---|---|---|---|---|---|---|---|---|---|---|---|
| **fixh** | * | 0.00 | -0.03 | 0.09 | 0.18 | -0.02 | -0.06 | 0.19 | -0.01 | -0.03 | 0.00 | -0.03 | 0.48 | -0.02 | -0.04 | -0.04 | 0.16 | 0.12 | -0.15 | -0.03 | -0.03 | -0.09 |
| **fum** | 0.00 | * | -0.03 | -0.01 | -0.04 | 0.00 | 0.29 | -0.09 | -0.03 | -0.03 | -0.05 | -0.05 | -0.03 | -0.03 | 0.01 | -0.04 | 0.00 | -0.05 | 0.04 | 0.04 | -0.03 | 0.09 |
| **holbo** | -0.03 | -0.03 | * | -0.09 | 0.19 | -0.05 | -0.05 | -0.08 | -0.16 | -0.03 | -0.08 | -0.04 | 0.05 | -0.02 | -0.03 | -0.04 | 0.24 | -0.04 | -0.12 | -0.06 | -0.03 | -0.15 |
| **mop** | 0.09 | -0.01 | -0.09 | * | 0.05 | -0.12 | 0.03 | 0.10 | 0.14 | 0.10 | -0.01 | -0.03 | 0.03 | -0.05 | -0.02 | -0.08 | 0.11 | 0.21 | 0.09 | 0.11 | -0.07 | 0.36 |
| **pa** | 0.18 | -0.04 | 0.19 | 0.05 | * | -0.11 | -0.06 | -0.06 | 0.00 | -0.01 | -0.16 | -0.10 | -0.06 | -0.05 | 0.27 | -0.07 | 0.38 | 0.13 | 0.06 | 0.11 | -0.03 | -0.07 |
| **rea** | -0.02 | 0.00 | -0.05 | -0.12 | -0.11 | * | 0.10 | 0.17 | 0.02 | 0.00 | 0.14 | 0.12 | -0.05 | -0.04 | -0.05 | -0.07 | -0.07 | -0.07 | 0.12 | -0.08 | 0.19 | -0.01 |
| **smo** | -0.06 | 0.29 | -0.05 | 0.03 | -0.06 | 0.10 | * | -0.08 | -0.08 | -0.05 | -0.05 | -0.05 | -0.05 | -0.03 | -0.06 | -0.07 | -0.08 | -0.07 | 0.04 | 0.09 | -0.02 | 0.03 |
| **stou** | 0.19 | -0.09 | -0.08 | 0.10 | -0.06 | 0.17 | -0.08 | * | 0.06 | -0.01 | 0.01 | 0.41 | 0.18 | 0.09 | -0.07 | 0.01 | -0.02 | 0.07 | -0.06 | -0.02 | -0.05 | -0.01 |
| **touo** | -0.01 | -0.03 | -0.16 | 0.14 | 0.00 | 0.02 | -0.08 | 0.06 | * | 0.02 | -0.08 | -0.08 | 0.06 | -0.05 | -0.03 | -0.11 | 0.02 | -0.03 | -0.02 | 0.17 | -0.07 | 0.04 |
| **area** | -0.03 | -0.03 | -0.03 | 0.10 | -0.01 | 0.00 | -0.05 | -0.01 | 0.02 | * | -0.02 | -0.05 | -0.03 | -0.02 | -0.04 | -0.03 | 0.10 | 0.09 | 0.10 | 0.01 | -0.02 | 0.10 |
| **talkagg** | 0.00 | -0.05 | -0.08 | -0.01 | -0.16 | 0.14 | -0.05 | 0.01 | -0.08 | -0.02 | * | 0.03 | 0.19 | -0.07 | -0.06 | -0.13 | -0.05 | -0.10 | -0.05 | -0.04 | 0.15 | 0.10 |
| **walkag** | -0.03 | -0.05 | -0.04 | -0.03 | -0.10 | 0.12 | -0.05 | 0.41 | -0.08 | -0.05 | 0.03 | * | -0.01 | -0.03 | -0.06 | -0.06 | -0.09 | -0.05 | -0.07 | -0.04 | -0.04 | 0.05 |
| **cry** | 0.48 | -0.03 | 0.05 | 0.03 | -0.06 | -0.05 | -0.05 | 0.18 | 0.06 | -0.03 | 0.19 | -0.01 | * | -0.02 | -0.03 | -0.03 | -0.05 | -0.04 | -0.07 | 0.03 | -0.03 | 0.00 |
| **leaf** | -0.02 | -0.03 | -0.02 | -0.05 | -0.05 | -0.04 | -0.03 | 0.09 | -0.05 | -0.02 | -0.07 | -0.03 | -0.02 | * | 0.01 | **0.65** | -0.02 | 0.00 | -0.10 | -0.03 | -0.02 | -0.01 |
| **scar** | -0.04 | 0.01 | -0.03 | -0.02 | 0.27 | -0.05 | -0.06 | -0.07 | -0.03 | -0.04 | -0.06 | -0.06 | -0.03 | 0.01 | * | 0.13 | 0.09 | -0.04 | -0.02 | 0.05 | -0.02 | -0.03 |
| **sitf** | -0.04 | -0.04 | -0.04 | -0.08 | -0.07 | -0.07 | -0.07 | 0.01 | -0.11 | -0.03 | -0.13 | -0.06 | -0.03 | 0.65 | 0.13 | * | -0.05 | 0.06 | -0.18 | -0.07 | -0.04 | 0.04 |
| **acr** | 0.16 | 0.00 | 0.24 | 0.11 | 0.38 | -0.07 | -0.08 | -0.02 | 0.02 | 0.10 | -0.05 | -0.09 | -0.05 | -0.02 | 0.09 | -0.05 | * | 0.11 | -0.10 | 0.07 | -0.05 | 0.07 |
| **handhip** | 0.12 | -0.05 | -0.04 | 0.21 | 0.13 | -0.07 | -0.07 | 0.07 | -0.03 | 0.09 | -0.10 | -0.05 | -0.04 | 0.00 | -0.04 | 0.06 | 0.11 | * | -0.15 | -0.01 | -0.03 | 0.08 |
| **handpo** | -0.15 | 0.04 | -0.12 | 0.09 | 0.06 | 0.12 | 0.04 | -0.06 | -0.02 | 0.10 | -0.05 | -0.07 | -0.07 | -0.10 | -0.02 | -0.18 | -0.10 | -0.15 | * | 0.15 | 0.15 | -0.11 |
| **lea** | -0.03 | 0.04 | -0.06 | 0.11 | 0.11 | -0.08 | 0.09 | -0.02 | 0.17 | 0.01 | -0.04 | -0.04 | 0.03 | -0.03 | 0.05 | -0.07 | 0.07 | -0.01 | 0.15 | * | -0.05 | -0.02 |
| **spit** | -0.03 | -0.03 | -0.03 | -0.07 | -0.03 | 0.19 | -0.02 | -0.05 | -0.07 | -0.02 | 0.15 | -0.04 | -0.03 | -0.02 | -0.02 | -0.04 | -0.05 | -0.03 | 0.15 | -0.05 | * | -0.04 |
| **talkg** | -0.09 | 0.09 | -0.15 | 0.36 | -0.07 | -0.01 | 0.03 | -0.01 | 0.04 | 0.10 | 0.10 | 0.05 | 0.00 | -0.01 | -0.03 | 0.04 | 0.07 | 0.08 | -0.11 | -0.02 | -0.04 | * |
