## Supplementary Table 3 for "An ethogram method for the analysis of human distress in the aftermath of public conflicts"

**Table S3.** Loadings of each behaviours on the five main principal components obtained from PCA. Different colours are indicative of the category to which each behaviour belongs.

| | *Components* | | | | |
|---|---|---|---|---|---|
| | *1* | *2* | *3* | *4* | *5* |
| fixh | 0.34 | 0.04 | **0.58** | -0.27 | 0.38 |
| fum | -0.08 | 0.15 | -0.21 | 0.19 | **0.52** |
| holbo | 0.31 | -0.02 | -0.15 | -0.47 | -0.05 |
| mop | 0.27 | 0.29 | 0.28 | **0.58** | -0.05 |
| pa | **0.65** | 0.21 | -0.14 | -0.24 | -0.16 |
| rea | -0.44 | 0.07 | 0.11 | -0.13 | -0.21 |
| smo | -0.21 | 0.16 | -0.23 | 0.17 | 0.49 |
| stou | -0.10 | -0.15 | **0.65** | 0.00 | -0.23 |
| touo | 0.07 | 0.23 | 0.15 | 0.27 | 0.01 |
| area | 0.10 | 0.15 | 0.02 | 0.26 | -0.27 |
| talkagg | -0.34 | 0.09 | 0.27 | -0.11 | 0.06 |
| walkag | -0.28 | -0.11 | 0.40 | -0.04 | -0.36 |
| cry | 0.07 | 0.02 | **0.59** | -0.25 | 0.48 |
| leaf | 0.11 | **-0.77** | -0.06 | 0.20 | 0.07 |
| scar | 0.30 | -0.07 | -0.23 | -0.08 | -0.07 |
| sitf | 0.15 | **-0.82** | -0.12 | 0.22 | 0.07 |
| acr | **0.62** | 0.18 | 0.01 | -0.11 | -0.15 |
| handhip | 0.38 | -0.02 | 0.20 | 0.25 | -0.17 |
| handpo | -0.24 | 0.39 | -0.26 | 0.04 | -0.16 |
| lea | 0.15 | 0.29 | -0.08 | 0.20 | 0.13 |
| spit | -0.29 | 0.08 | -0.08 | -0.19 | -0.17 |
| talkg | 0.03 | 0.08 | 0.18 | **0.62** | -0.02 |
