## Supplementary Table 4 for "An ethogram method for the analysis of human distress in the aftermath of public conflicts"

**Table S4.** Eigenvalues, proportion of variation explained and cumulative percentage explained of each dimension (Dim.) obtained from the PCA.

| *Dimension* | *Eigenvalue* | *Proportion of variation explained* | *Cumulative percentage explained* |
|---|---|---|---|
| Dim. 1 | 1.98 | 9 | 9 |
| Dim. 2 | 1.86 | 8.46 | 17.46 |
| Dim. 3 | 1.81 | 8.24 | 25.71 |
| Dim. 4 | 1.62 | 7.35 | 33.05 |
| Dim. 5 | 1.35 | 6.16 | 39.21 |
| Dim. 6 | 1.29 | 5.87 | 45.08 |
| Dim. 7 | 1.27 | 5.76 | 50.84 |
| Dim. 8 | 1.19 | 5.39 | 56.23 |
| Dim. 9 | 1.09 | 4.94 | 61.17 |
| Dim. 10 | 1 | 4.54 | 65.71 |
| Dim. 11 | 0.96 | 4.37 | 70.07 |
| Dim. 12 | 0.91 | 4.14 | 74.21 |
| Dim. 13 | 0.83 | 3.76 | 77.97 |
| Dim. 14 | 0.75 | 3.42 | 81.39 |
| Dim. 15 | 0.71 | 3.21 | 84.6 |
| Dim. 16 | 0.67 | 3.05 | 87.64 |
| Dim. 17 | 0.61 | 2.79 | 90.43 |
| Dim. 18 | 0.55 | 2.5 | 92.93 |
| Dim. 19 | 0.48 | 2.18 | 95.11 |
| Dim. 20 | 0.46 | 2.09 | 97.2 |
| Dim. 21 | 0.31 | 1.4 | 98.6 |
| Dim. 22 | 0.31 | 1.4 | 100 |
